# Lipooligosaccharide, Vag8, and pertussis toxin of *Bordetella pertussis* cooperatively cause coughing in mice

**DOI:** 10.1101/2020.12.25.424096

**Authors:** Yukihiro Hiramatsu, Koichiro Suzuki, Takashi Nishida, Naoki Onoda, Takashi Satoh, Shizuo Akira, Masahito Ikawa, Hiroko Ikeda, Junzo Kamei, Sandra Derouiche, Makoto Tominaga, Yasuhiko Horiguchi

## Abstract

Whooping cough, a contagious respiratory disease caused by *Bordetella pertussis*, is characterized by paroxysmal coughing; however, the mechanism has not been studied because of the lack of versatile animal models that reproduce the cough. Here, we present a mouse model that reproduces coughing after intranasal inoculation with the bacteria or its components and demonstrate that lipooligosaccharide (LOS), pertussis toxin (PTx), and Vag8 of the bacteria cooperatively function to cause coughing. LOS-induced bradykinin sensitized a transient receptor potential ion channel, TRPV1, which acts as a sensor to evoke the cough reflex. Vag8 further increased bradykinin levels by inhibiting the C1 esterase inhibitor, the major downregulator of the contact system, which generates bradykinin. PTx inhibits intrinsic negative regulation systems for TRPV1 through inactivation of G_i_ GTPases. Our findings provide a basis for answering long-standing questions on the pathophysiology of the pertussis cough.

## Introduction

Whooping cough, also referred to as pertussis, is a highly contagious respiratory disease caused by the gram-negative bacterium *Bordetella pertussis* (Kilgore et al., 2016). Generally, the disease can be prevented through vaccination; however, the number of pertussis cases is significantly increasing worldwide, hypothetically because of the rapid waning of immunity induced by recent acellular vaccines and adaptation of the bacteria to escape vaccine-induced immunity(Tan et al., 2015; Twillert et al., 2015). Patients with the disease exhibit various clinical manifestations, including bronchopneumonia, pulmonary hypertension, hypoglycemia, leukocytosis, and paroxysmal coughing. Among these, paroxysmal coughing, which persists for several weeks and imposes a significant burden on infants, is the hallmark of pertussis. However, etiological agents and the mechanism of pertussis cough remain unknown.

*B. pertussis* is highly adapted to humans; therefore, convenient animal models that reproduce coughing induced by the bacterial infection are difficult to establish. Baboons were recently reported to replicate various pertussis symptoms, including paroxysmal coughing (Warfel et al., 2012a; 2012b; Warfel and Merkel, 2014); however, these large primates are difficult to use in a sufficient number for analytical experiments because of ethical and cost issues. Rats were reported to cough after intrabronchial administration of agarose-embedded or naked *B. pertussis* (Hall et al., 1999; 1998; 1997; 1994; Hornibrook and Ashburn, 1939; Parton et al., 1994; Wardlaw et al., 1993; Woods et al., 1989); however, there have been no reports on further studies with rats that analyze the mechanism underlying the coughing. Meanwhile, we re-established the coughing model of rats infected with *B. bronchiseptica*, which produces many common virulence factors shared with *B. pertussis*, and observed that BspR/BtrA, an anti-σ factor, regulates the ability of *B. bronchiseptica* to cause coughing in rats (Nakamura et al., 2019). Additionally, our rat model exhibited coughing in response to *B. pertussis* infection; however, the cough frequencies were lower than that caused by *B. bronchiseptica*, and the cough production was not reproduced well. Therefore, we attempted to develop an alternative animal model for pertussis cough and focused on mice because of their multiple genetically modified mutants. Mice have not been used for analyses of *B. pertussis-induced* coughing, because they have long been believed to be unable to cough (Elahi et al., 2007; Mackenzie et al., 2004; Scanlon et al., 2019). However, recently, many reports have demonstrated that vagal sensory neurons, which are involved in the cough reflex, innervate the mouse airway (Dinh et al., 2005; Thai Dinh et al., 2005; J. W. Zhang et al., 2006). Mouse coughing in response to certain tussive stimuli has been detected and recorded by independent research groups (Chen et al., 2013; Kamei et al., 1993; 2006; Lin et al., 2019; C. Zhang et al., 2017). In the present study, our mouse-coughing model revealed that lipooligosaccharide (LOS), Vag8, and pertussis toxin (PTx) cooperatively function to produce coughing through the pathway from bradykinin (Bdk) generation to transient receptor potential vanilloid 1 (TRPV1) sensitization.

## Results

### C57BL/6 mice respond to *B. pertussis* infection by coughing

We intranasally inoculated mice with *B. pertussis* in a manner similar to that in our previous study on rats (Nakamura et al., 2019) and observed that C57BL/6J mice coughed approximately one week after inoculation (Movies S1 and S2, and Fig. 1A–1C). The incidence of coughing varied depending on the bacterial strains and mouse strains. *B. pertussis* 18323, a laboratory strain, markedly caused coughing while another classical strain, Tohama, did not. Clinical isolates caused varying degrees of coughing. This effect did not correlate with their ability to colonize the trachea and lungs. Unlike C57BL/6J mice, BALB/c mice did not exhibit coughing upon infection with *B. pertussis* 18323 (Fig. S1A–S1B). The extent of cough production did not differ between the sexes of the mice (data not shown). Thus, we further explored the mechanism of pertussis cough using *B. pertussis* 18323 and male C57BL/6J mice.

**Fig. 1.**
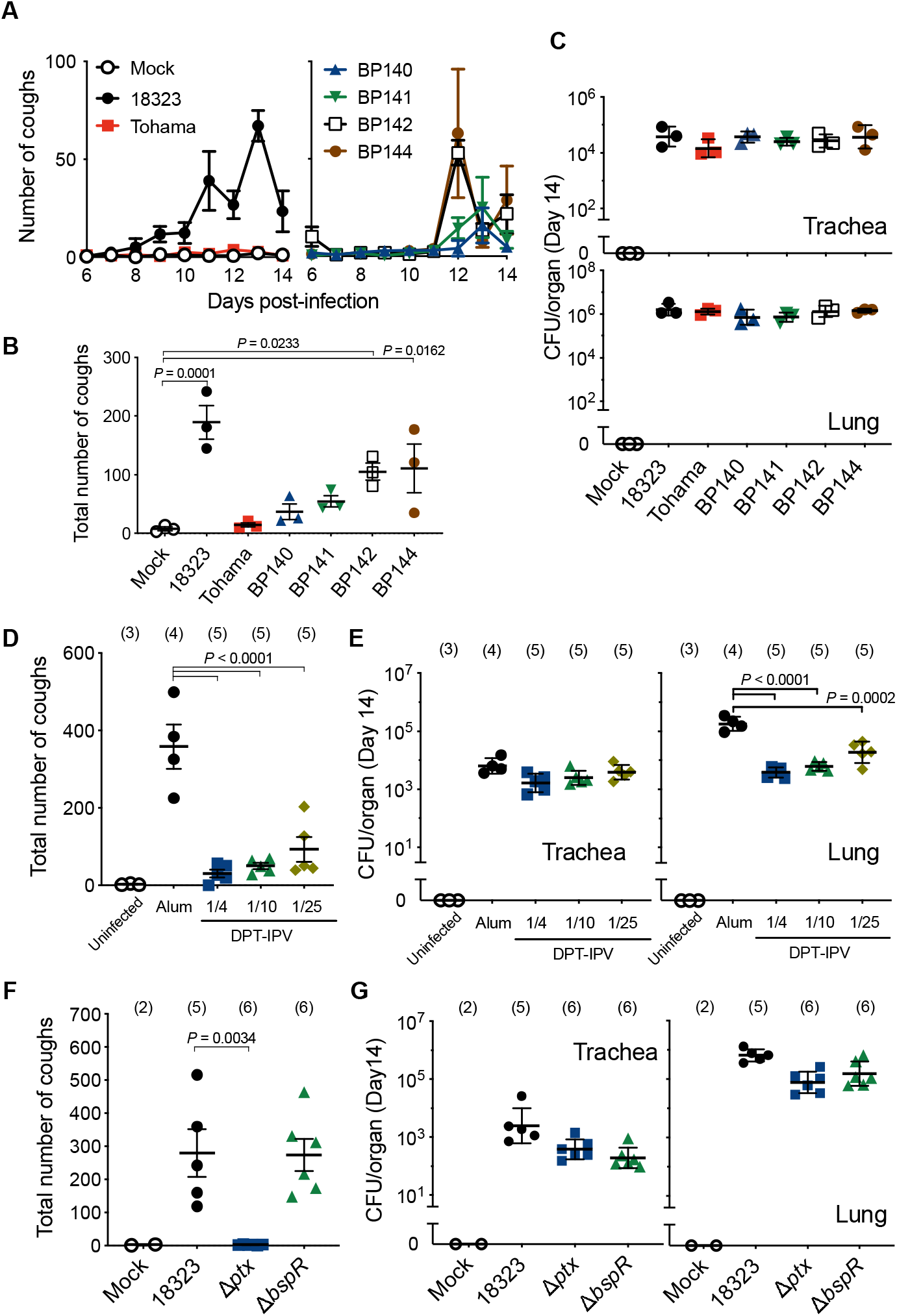
Coughing of mice inoculated with *B. pertussis*. (**A–C**) Cough production in male C57BL/6J mice inoculated with *B. pertussis* 18323, Tohama, BP140, BP141, BP142, or BP144 (n = 3). (**D** and **E**) Cough production in mice immunized with DPT-IPV. Male C57BL/6J mice were immunized with 1/4, 1/10, or 1/25 the human dose of DPT-IPV or 4 mg of aluminum hydroxide (Alum). (**F** and **G**) Cough production in male C57BL/6J mice inoculated with wild-type (18323), Δ*ptx*, or Δ*bspR* strains of *B. pertussis*. Mice were intranasally inoculated with 5 × 10^6^ CFU of *B. pertussis* strains in 50 μl of SS medium (Mock). The number of coughs was counted for 5 min/mouse/day for 9 days from days 6 to 14 post-inoculation (A), and the total number is expressed (B, D, and F). The number of bacteria recovered from the tracheas and lungs was counted on day 14 (C, E, and G). Each plot represents the mean ± SEM (A). Each horizontal bar represents the mean ± SEM (B, D, and F) or geometric mean ± SD (C, E, and G). The number of mice in each test group is presented in parentheses (D–G).

### Pertussis toxin (PTx) contributes to cough production

Previous studies using rat or baboon models of *B. pertussis* infection suggested the involvement of PTx in cough production based on the observation that infection with a PTx-deficient strain did not cause coughing and that immunization with acellular pertussis vaccines containing pertussis toxoid protected the animals from coughing after the bacterial infection (Hall et al., 1998; Parton et al., 1994; Warfel et al., 2013). Therefore, we examined this idea by immunizing mice with the DPT-IPV (diphtheria-pertussis-tetanus-inactivated polio) vaccine. Consistent with previous studies, the immunized mice were colonized by the bacteria but exhibited no or less coughing (Fig. 1D, 1E, S1C, and S1D). A PTx-deficient strain (Δ*ptx*) did not cause coughing in mice (Fig. 1F and 1G). A mutant strain of *B. pertussis* that lacked *bspR*, a transcriptional regulator associated with the cough-inducing ability of *B. bronchiseptica* in rats (Nakamura et al., 2019), caused coughing. This result indicates that the regulatory systems of bacterial gene expression that cause coughing differ between *B. pertussis* and *B. bronchiseptica*. We previously observed that not only living *B. bronchiseptica* but also the bacterial lysates caused coughing in rats (Nakamura et al., 2019). Similarly, in the present study, the lysates of wild-type *B. pertussis* caused coughing similar to the infection with living bacteria (Fig. 2A and 2B). In contrast, the lysates from neither the Δ*Rptx* mutant nor the mutant producing enzymatically inactive PTx (PTx_ED_) caused coughing (Fig. 2A, 2B and S2A). The Δ*ptx* lysate complemented with purified PTx caused coughing to the same extent as the wild-type lysate (Fig. 2B and 2D). These results indicate that the enzyme (ADP-ribosylating) activity of PTx is required for cough production. However, contrary to our expectation, intranasal inoculation of purified PTx hardly caused coughing, implying that bacterial factors other than PTx are required (Fig. 2A and 2B). The involvement of FhaB, which is also included in the DPT-IPV vaccine, was concluded to be negligible because the lysate of the FhaB-deficient mutant caused coughing as well (Fig. 2C).

**Fig. 2.**
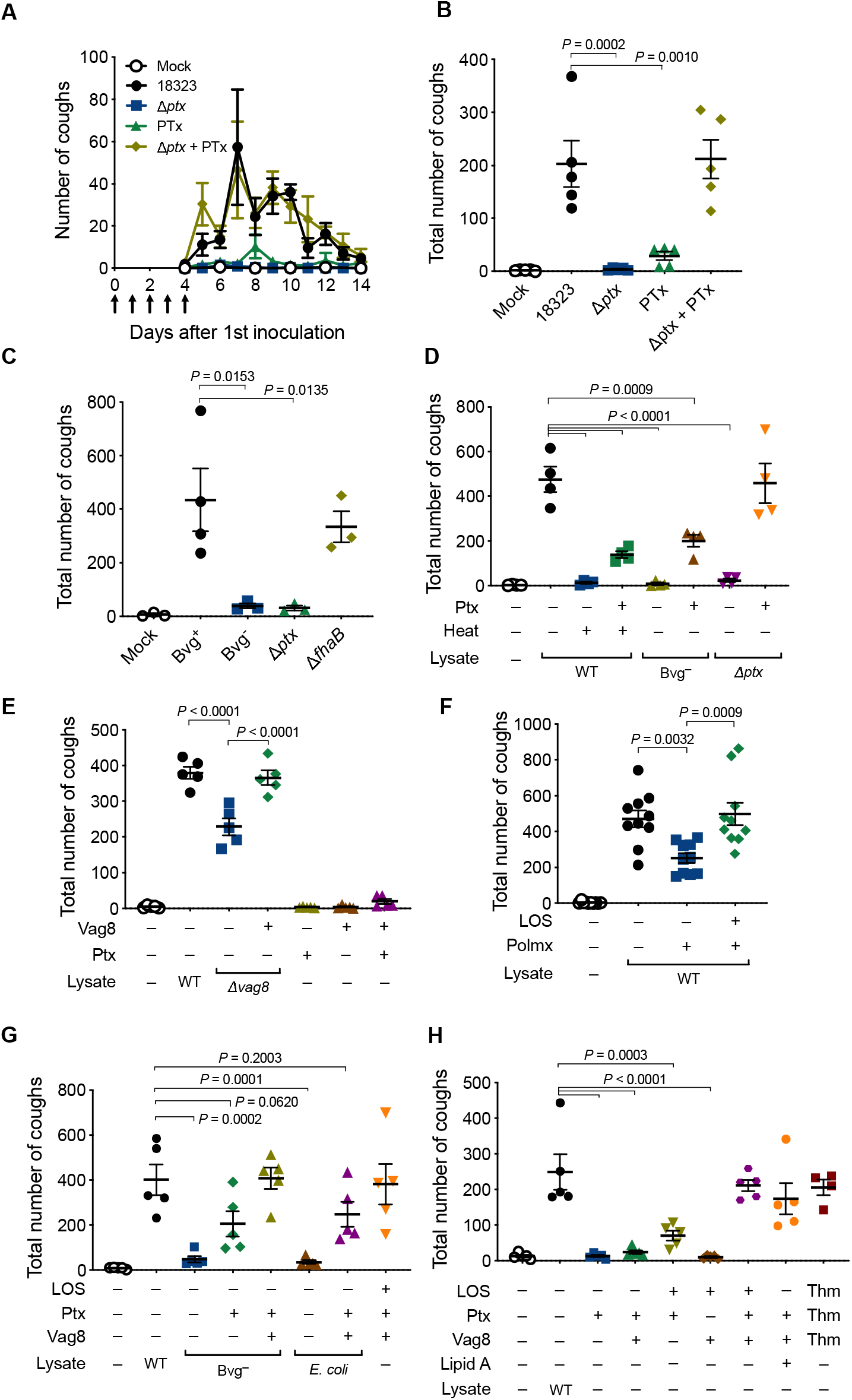
Coughing of mice inoculated with *B. pertussis* components. (**A–C**) Cough production in mice inoculated with cell lysates from *B. pertussis* wild-type and mutant strains. Mice were intranasally inoculated with cell lysates (50 μg) of *B. pertussis* 18323 wild-type, Δ*ptx* strains, or PTx (200 ng) with or without Δ*ptx* cell lysate on days 0 to 4 (arrows in A). The number of coughs was counted for 11 days from days 4 to 14 (A) and expressed as the sum from days 6 to 14 (B). Mice were similarly inoculated with cell lysates of *B. pertussis* wild-type, Bvg^+^-phase locked, Bvg^−^-phase locked, Δ*ptx*, or Δ*fhaB* strains. The total number of coughs from days 6 to 14 is shown (C). (**D–H**) Cough production in mice inoculated with various preparations of bacterial components. The number of coughs was counted, and the sum of coughs from days 6 to 14 is shown. The inoculated preparations were as follows: cell lysates (50 μg) from *B. pertussis* 18323 wild-type (WT) (D-H), Δ*ptx* (D), Bvg^−^-phase locked (D and G), Δ*vag8* (E), and *E. coli* DH5α (G); cell lysates of *B. pertussis* 18323 wild-type that were incubated at 56°C for 1 h (D, Heat +) or pretreated with Detoxi-Gel™ (polymyxin B, Polmx +) (F); PTx (D, E, G, and H, 200 ng), Vag8 (E, G, and H, 500 ng), and LOS (F–H, 4 × 10^4^ EU) of the 18323 strain; and synthetic lipid A of *E. coli* (H, 4 × 10^4^ EU). PTx, Vag8, and LOS of the Tohama strain are indicated as “Thm”. PBS was used for mock-inoculation. Each plot represents the mean ± SEM (A). Each horizontal bar represents the mean ± SEM (B–H). The number of mice in each test group is as follows: n = 5 (A and B), n = 3 or 4 (C), n =4 (D), n = 5 (E and G), n = 10 (F), n = 4 or 5 (H).

### Vag8 and lipooligosaccharide (LOS) along with PTx contribute to cough production

To identify additional bacterial factors contributing to cough production, we examined various combinations of bacterial lysates and purified bacterial components. *B. pertussis* exhibits two distinct phenotypic phases, Bvg^+^ and Bvg^-^, in response to environmental alterations (Cotter and Jones, 2003). In the Bvg^+^ phase, the bacteria produce a set of virulence factors including PTx, adenylate cyclase toxin, dermonecrotic toxin, and the machinery and effectors of the type III secretion system. In contrast, in the Bvg^-^ phase, the bacteria shut down the expression of the virulence factors and, instead, express several factors specific to this phenotype. Therefore, in general, the Bvg^+^ phase is considered to represent the virulent phenotype of the organism. Consistently, the lysate from the Bvg^+^ phase-locked mutant (Bvg^+^ lysate) caused coughing, while that from the Bvg^-^ phase-locked mutant (Bvg^-^ lysate) did not (Fig. 2C). The addition of PTx into the Bvg^-^ lysate only partially restored the cough production, compared to the wild-type lysate and the Δ*px* lysate complemented with PTx (Fig. 2D). The wild-type lysate and Δ*ptx* lysate were obtained from the bacteria in the Bvg^+^ phase. Therefore, these results suggest that at least two distinct factors in addition to PTx are involved in cough production. One factor is present in the Bvg^+^ lysate, and the other is a Bvg-independent molecule present both in the Bvg^+^ and Bvg^-^ lysates. Heat treatment at 56°C for 1 h abrogated the cough induced by the wild-type lysate (Fig. 2D). The addition of PTx to the heat-treated lysate moderately restored the cough production, compared to the wild-type lysate and the Δ*ptx* lysate complemented with PTx. Considering that purified PTx alone did not cause coughing, we hypothesize that one factor is heat-labile and the other factor is heat-stable.

We next examined various mutants of *B. pertussis* kept in our laboratory and observed that a mutant strain (Δ*vag8*) that is deficient in the autotransporter protein Vag8 exhibited only a moderate ability to cause coughing, whereas another autotransporter mutant, Δ*brkA*, was fully active (Fig. S2B and S2C). Similarly, the Δ*vag8* lysate caused moderate coughing compared to the wild-type lysate (Fig. 2E). The addition of a recombinant Vag8 protein (Vag8p) compensated for the activity of the Δ*vag8* lysate. Vag8p alone or in combination with PTx did not cause coughing (Fig. 2E). The Bvg^-^ lysate complemented with PTx and Vag8p caused coughing to the same extent as the wild-type lysate (Fig. 2G). These results revealed that Vag8 is the second factor contributing to cough production. Because Vag8 is heat-labile and specific to the Bvg^+^-phase bacteria, we narrowed the third factor to one that is heat-stable and present independently of the Bvg phases. We considered LOS, which is a heat-stable and biologically active outer membrane component, as a probable candidate. The wild-type lysate in which LOS was eliminated with polymyxin B treatment exhibited a reduced ability to cause coughing (Fig. 2F). The addition of a purified LOS preparation restored the coughing. When the lysate from *E. coli* DH5α was inoculated, instead of that from *B. pertussis* in combination with PTx and Vag8, the mice exhibited coughing (Fig. 2G). Finally, the combination of PTx, Vag8, and LOS caused coughing to the same extent as the wild-type lysate (Fig. 2G and 2H). Thus, we concluded that LOS, PTx, and Vag8 cooperatively cause coughing in mice; however, the combination of any two of these three factors caused no or only a few coughs (Fig. 2H). Notably, commercially available synthetic lipid A of *E. coli* could serve as a substitute for LOS for producing coughing in combination with PTx and Vag8 (Fig. 2H). These results indicate that the lipid A moiety of LOS, which is virtually identical among gram-negative bacteria, is essential to cause coughing. LOS, PTx, and Vag8 from the *B. pertussis* Tohama strain caused coughing in mice, while the living bacteria or the lysate of this strain did not (Fig. 1A, 1B, and 2H). PTx from the Tohama strain compensated for the ability of the Δ*ptx* lysate of the 18323 strain to cause coughing (Fig. S2D). In contrast, PTx from both Tohama and 18323 did not compensate the ability of the Δ*ptx* lysate of Tohama strain. These results suggest that the Tohama strain may produce an inhibitory factor or multiple inhibitory factors against cough production.

### Cough-evoking pathways stimulated by LOS, Vag8, and PTx

The cough reflex involves vagal afferent fibers innervating the airways from the larynx to the proximal bronchi; however, the mechanism through which the afferent nerves are activated to evoke coughing is not fully understood. Nevertheless, it is accepted that transient receptor potential (TRP) ion channels on sensory nerve terminals participate in the cough reflex (Bonvini and Belvisi, 2017; Grace et al., 2011). TRP channels are modulated by cell signals from G protein-coupled receptors (GPCRs) that are activated by ligands such as Bdk, prostanoids, and tachykinins including neurokinins and substance P (Bonvini and Belvisi, 2017; Carr et al., 2003; Geppetti et al., 2009; Grace et al., 2012; 2011; Kumar et al., 2017; Maher et al., 2012; 2011; Mizumura et al., 2009; Sugiura et al., 2002). To explore the stimulatory mechanism through which the *B. pertussis* components cause coughing, we utilized various antagonists for GPCRs (Bdk B1 and B2 receptors, Neurokinin 1–3 receptors, and prostaglandin E2 receptor EP3) and TRP channels (TRPV1, TRPV4, and TRPA1) in our mouse model of coughing. Mice pre-administered antagonists against the B2 receptor (B2R) and TRPV1 exhibited reduced levels of coughing in response to bacterial lysate; however, other reagents were apparently ineffective (Fig. 3A). Similar results were obtained in mice inoculated with LOS, PTx, and Vag8 (Fig. 3B, S3A, and S3B). When both antagonists against B2R and TRPV1 were simultaneously administered, the inhibitory effects were not additively augmented, suggesting that the pathway upstream of B2R to TRPV1 may neither diverge nor converge (Fig. S3C). TRPV1-deficient mice, but not TRPA1-deficient mice, were less responsive to the combination of LOS, PTx, and Vag8 (Fig. 3C); this observation is consistent with the results of the experiments using the antagonists. Thus, we considered LOS, PTx, and Vag8 to cooperatively stimulate the afferent nerves through a pathway from Bdk–B2R to TRPV1 (Fig. 6E). Bdk was reported to enhance the cough reflex in guinea pigs (Al-Shamlan and El-Hashim, 2019; Featherstone et al., 1996; Fox et al., 1996). Bdk concentrations in the bronchoalveolar lavage fluid (BALF) of mice were increased 4 days after intranasal inoculation with LOS–PTx–Vag8 (Fig. 3D). We further analyzed the roles of each bacterial component in the stimulating pathway from Bdk to TRPV1.

**Fig. 3.**
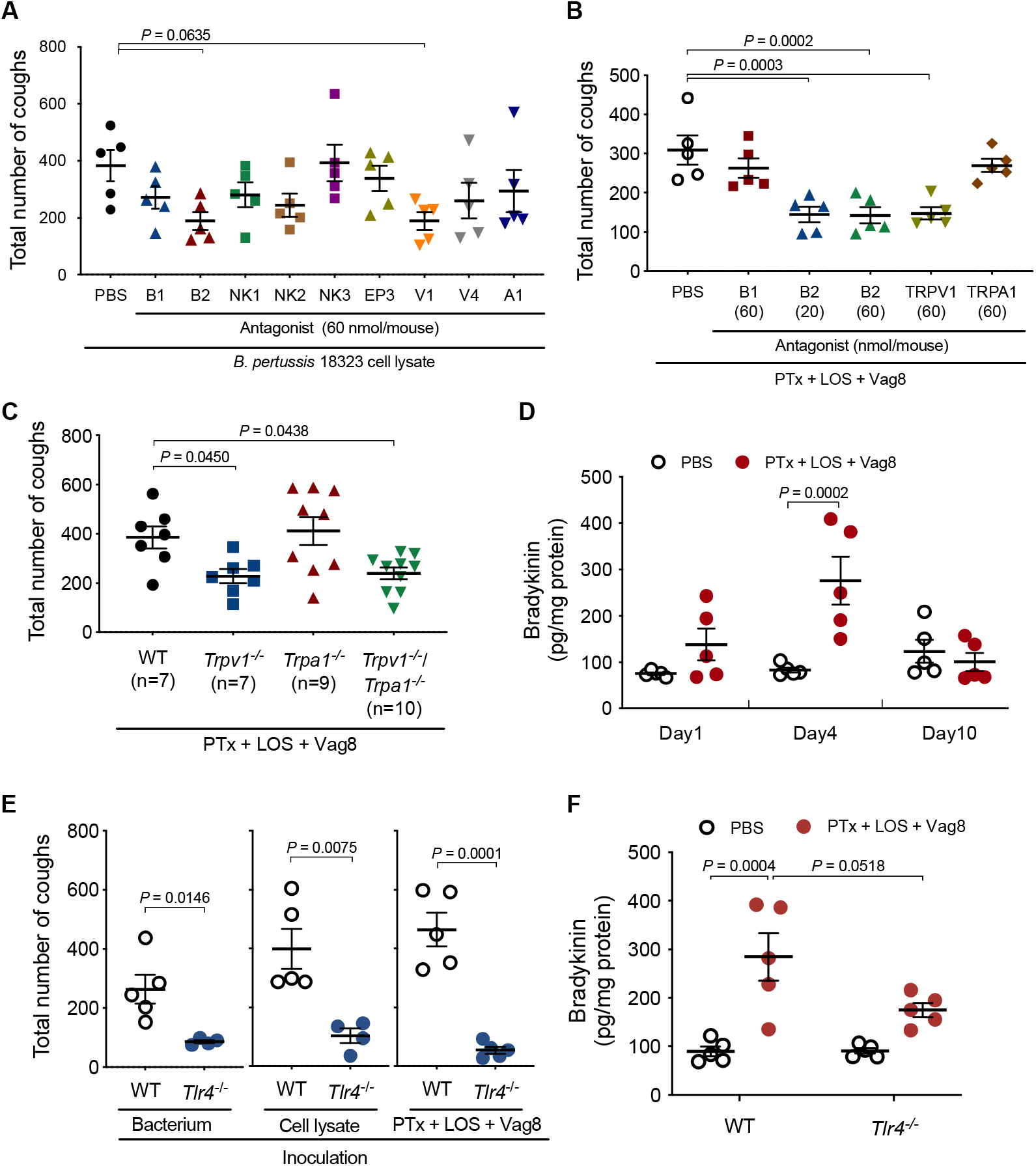
Involvement of B2R, TRPV1, and TLR4 in *B. pertussis-induced* coughing. (**A** and **B**) Effects of antagonists on cough reflex-related pathways in *B. pertussis-induced* coughing. Mice were inoculated with each antagonist prior to inoculation with *B. pertussis* 18323 cell lysate (A, 50 μg) or combinations (B) of PTx (200 ng), LOS (4 × 10^4^ EU), and Vag8 (500 ng). The number of coughs was counted as described in Materials and Methods. The sum of coughs from days 6 to 14 is shown. V1, TRPV1; V4, TRPV4; A1, TRPA1 (A). (**C** and **E**) Cough production of mice deficient in TRP ion channels (C) or TLR4 (E). Wild-type (WT), *Trpv1*^-/-^, *Trpa1*^-/-^, *Trpv1*^-/-^/ Trpa1^-/-^, or *Tlr4*^-/-^ mice were inoculated with *B. pertussis* 18323 (F, left panel), the cell lysate of the bacteria (F, center panel), or the combination (C and F, right panel) of PTx, LOS, and Vag8. The sum of coughs from days 6 to 14 is shown. (**D** and **F**) Bdk concentrations in the BALF of mice inoculated with the combination of PTx, LOS, and Vag8. Wild-type (WT) and *Tlr4*^-/-^ mice were similarly inoculated with the sample, and the concentrations of Bdk in BALF on days 1, 4, and 10 (D) or 4 (F) were determined using ELISA. Each horizonal bar represents the mean ± SEM. The number of mice in each test group is as follows: n = 5 (A, B, D, and F), n = 7 to 10 (C), n = 4 or 5 (E). Two-way ANOVA with (F) or without (D) Sidak’s multiple-comparison test or unpaired *t*-test (E) was performed for the statistical analyses.

### Role of LOS

Because the lipid A moiety of LOS was determined to be essential for cough production, we examined the role of the specific receptor for lipid A, toll-like receptor 4 (TLR4), by using TLR4-deficient mice. *B. pertussis* colonized the respiratory organs of TLR4-deficient mice, similar to that in isogenic wild-type mice (Fig. S4A); however, it caused low cough production in the deficient mice (Fig. 3E). Similar results were obtained with intranasal inoculation of the mice with the bacterial lysates or the combination of LOS–PTx–Vag8 (Fig. 3E). The increase in Bdk levels in the BALF of TLR4-deficient mice inoculated with LOS–PTx–Vag8 was considerably lower than that of wild-type mice (Fig. 3F). These results suggest that LOS stimulates Bdk generation via interaction with TLR4. In addition to Bdk, proinflammatory cytokines such as IL-1β, IL-6, and TNFα tended to increase in wild-type mice but not in TLR4-deficient mice inoculated with LOS–PTx–Vag8 (Fig. S4B). Furthermore, increases in these cytokines were not observed without inoculation of LOS (Fig. S4C), indicating that LOS leads to slight inflammation via TLR4 under these experimental conditions. Bdk is a potent inflammatory mediator that is released from high-molecular-weight kininogen (HK) through the proteolytic activity of plasma kallikrein (PK). HK-deficient (*Kng1*^-/-^) mice, which do not produce Bdk, were less responsive to infection with *B. pertussis* or inoculation of the bacterial lysate than isogenic wild-type mice (Fig. 4A and 4B), demonstrating that Bdk participates in a cascade leading to *B. pertussis-induced* cough.

**Fig. 4.**
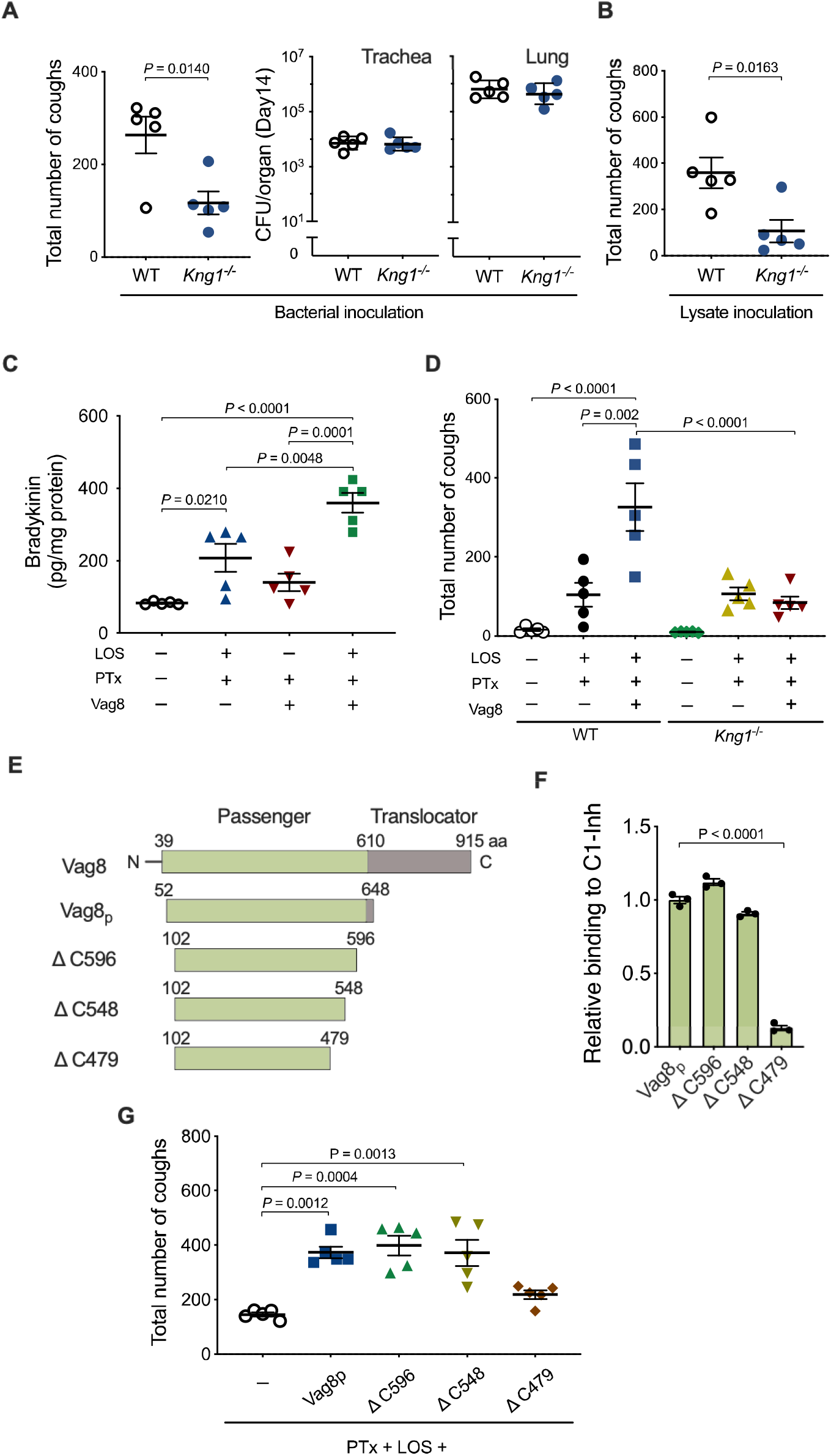
Role of Vag8 in cough production. (**A, B**, and **D**) Cough production of *Kng1*^-/-^ mice. Wild-type (WT) and *Kng1*^-/-^ mice were inoculated with *B. pertussis* 18323 (A); the cell lysate of the bacteria (B); or combinations (D) of PTx, LOS, and Vag8. The sum of coughs from days 6 to 14 (A left panel, B, and D) and the number of the bacteria recovered from the trachea and lungs on day 14 (A right panel) are shown. (**C**) Bdk concentrations in the BALF of mice inoculated with PTx, LOS, and/or Vag8. (**E**) Schematic representations of Vag8 and recombinant proteins. The passenger domain of Vag8 (Vag8_p_) and truncated derivatives are listed with their names and amino-acid positions. (**F**) The binding of the Vag8 recombinant proteins to C1-Inh. Relative binding levels of the Vag8 recombinant proteins are expressed as OD_450_ values normalized to those for Vag8_p_ in the ELISA-based binding assay. Bars represent the means and SEM (n = 3). (**G**) Cough production of mice inoculated with Vag8_p_ (500 ng, ca. 8 pmol) or the truncated derivatives (ΔC596, ΔC548, or ΔC479, 24 pmol) along with PTx and LOS. The sum of coughs from days 6 to 14 are shown. Each horizonal bar represents the mean ± SEM (A left panel, B–D, and G) or geometric mean ± SD (A right panel). Unpaired *t*-test was performed for the statistical analyses (A and B).

### Role of Vag8

Bdk, HK, and PK comprise the kallikrein–kinin system in the plasma contact system, which is initiated and accelerated by two proteases, factor XII (FXII) and plasma prekallikrein (PPK) (Maas et al., 2011; Weidmann et al., 2017; Wu, 2018) (Fig. 6E). Certain stimuli convert zymogen FXII into an active enzyme, FXIIa. FXIIa cleaves PPK to generate PK, which releases Bdk by cleaving HK. In turn, PK cleaves FXII to release FXIIa, providing a positive feedback cycle. FXIIa and PK are inhibited by C1 esterase inhibitor (C1-Inh), the major regulator of the contact system, which is present in the human plasma (Sanrattana et al., 2019). This inhibitory action of C1-Inh was reportedly inhibited by Vag8 (Hovingh et al., 2017; Marr et al., 2011). If this is the case, Vag8 is likely to exacerbate the cough response of mice by upregulating the Bdk level through the inhibition of C1-Inh activity. Indeed, the Bdk levels in the BALF of mice inoculated with LOS and PTx were increased upon the administration of Vag8p (Fig. 4C). These results are consistent with the results described above showing that Vag8p increased the number of coughs induced by the Δ*vag8* lysate or the combination of LOS and PTx (Fig. 2E, 2H, and 4D). This exacerbating effect of Vag8p on cough production was not observed in *Kng1*^-/-^ mice (Fig. 4D). In addition, the truncated fragment of Vag8p ranging from amino acid positions 102 to 479 (ΔC479), which is unable to bind and inactivate C1-Inh, did not increase the number of coughs in mice inoculated with LOS and PTx; in contrast, other types of truncated Vag8, which retain the ability to inhibit C1-Inh increased the number of coughs similar to the whole passenger-domain fragment of Vag8 (Vag8p) (Onoda et al., 2020) (Fig. 4E–4G). These results indicate that Vag8 exacerbates the cough response of mice by upregulating the Bdk level through the inhibition of C1-Inh activity.

### Role of PTx in Bdk-induced sensitization of TRPV1

The above results indicate that LOS and Vag8 function to raise the level of Bdk. Bdk sensitizes TRPV1, which is directly activated by capsaicin (Cap) to evoke coughing in animals (Bonvini and Belvisi, 2017; Caterina et al., 1997; Choi and Hwang, 2018; Fox et al., 1996; Grace et al., 2012; Shin et al., 2002; Trevisani et al., 2004). Previous reports indicated that PTx enhanced the Bdk action by affecting the GTPases of the G_i_ family, which are the target molecules of the toxin (Diehl et al., 2014; Moss et al., 1988). We therefore examined the role of PTx in the effect of Bdk on TRPV1 using a whole-cell patch clamp technique in HEK293T cells expressing B2R and TRPV1. The cells without ectopic expression did not respond to Bdk and Cap (data not shown). Cap induced a TRPV1-dependent current that was desensitized in the second Cap application (Fig. 5A and 5B) (Bhave et al., 2002; Mohapatra, 2003; Mohapatra and Nau, 2005; Sanz-Salvador et al., 2012). When the cells were treated with Bdk before the second application, the Cap-induced currents were not reduced but were enhanced in the second application, as reported previously (Fox et al., 1996). We observed that this enhancement was further augmented in cells pretreated with PTx but not in those pretreated with PTx_ED_. Without Bdk, PTx did not influence the desensitization induced by the second Cap application. Similar results were obtained in independent experiments that evaluated the intracellular Ca^2+^ level of dorsal root ganglion (DRG) neurons (Fig. 5D).

**Fig. 5.**
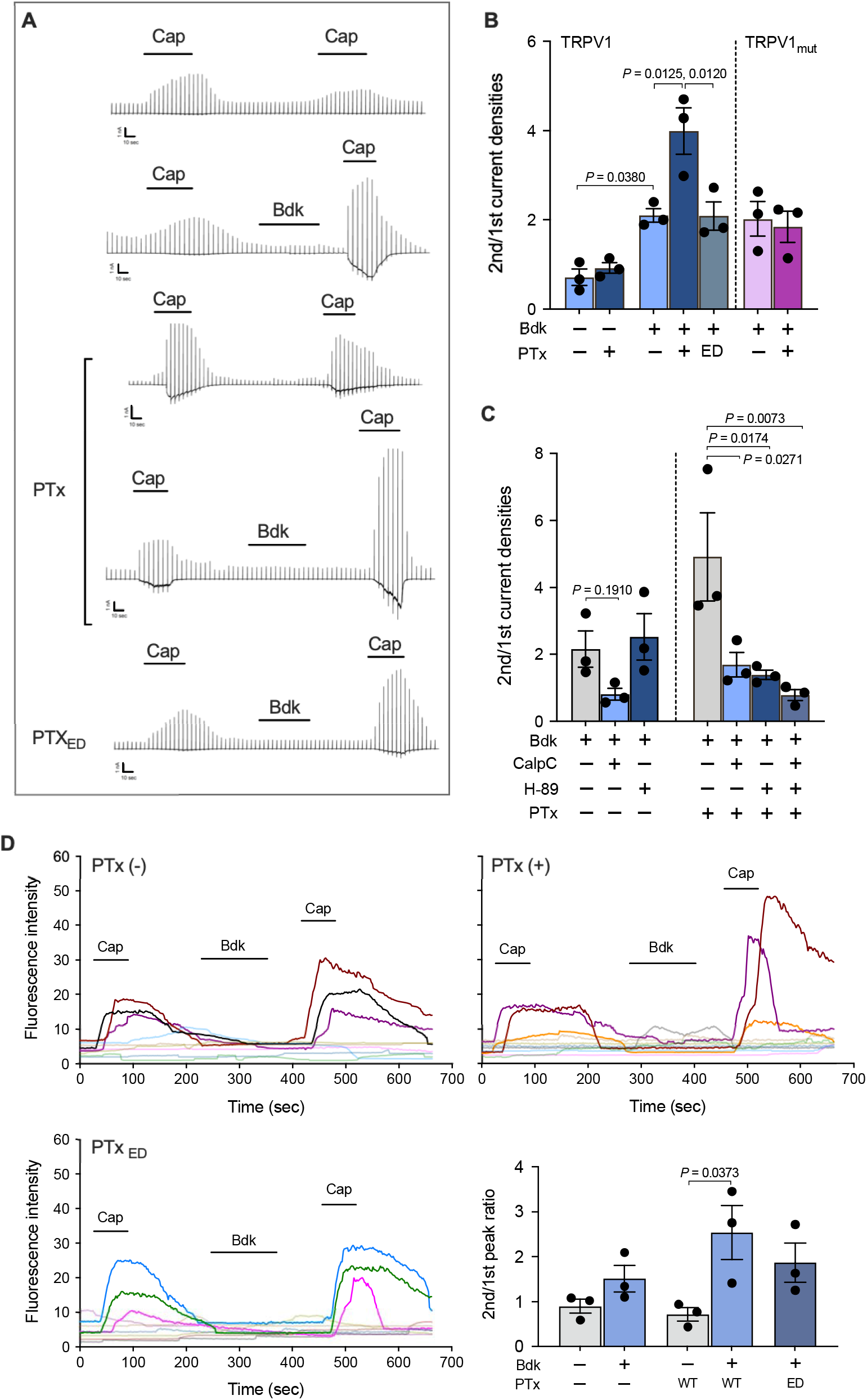
Role of PTx in Bdk-induced sensitization of TRPV1. (**A**) Current responses of HEK293T cells expressing B2R and wild-type TRPV1 or mutant (TRPV1_mut_) to repetitive applications of Cap (10 nM) and an intervening application of Bdk (100 nM). Representative traces of Cap-evoked currents are shown. Scales at the left bottom of each trace indicate 1 nA and 10 sec on the ordinate and abscissa, respectively. Horizontal bars below the reagent names indicate the incubation period of each reagent. (**B** and **C**) Ratio of the second peak to the initial peak of the current densities induced by Cap. The cells were preincubated with or without 10 ng/ml PTx or the enzymatically inactive derivative of PTx (PTx_ED_) for 24-30 h prior to the recording (A–C). The cells were stimulated with Bdk and subsequently Cap in the presence of 1 μM CalpC and/or H-89 (C). Values represent the means ± SEM (n = 3). See also Fig. S5. (**D**) Intracellular calcium levels of DRG cells changed in response to repetitive applications of Cap. Isolated DRG cells pretreated with or without 10 ng/ml PTx or PTx_ED_ for 24–30 h were subjected to calcium imaging with transient applications of Cap (1 μM for 1 min) and Bdk (100 nM for 3 min). Each line represents a single cell isolated from DRG. Nine to ten independent cells were tested for each experiment. The results for Cap-sensitive neurons are strongly colored (upper and lower left panels). (Lower right panel) Fluorescence-intensity ratios of the second peak to the initial peak increased in response to Cap. Values represent the means ± SEM (n = 3).

**Fig. 6.**
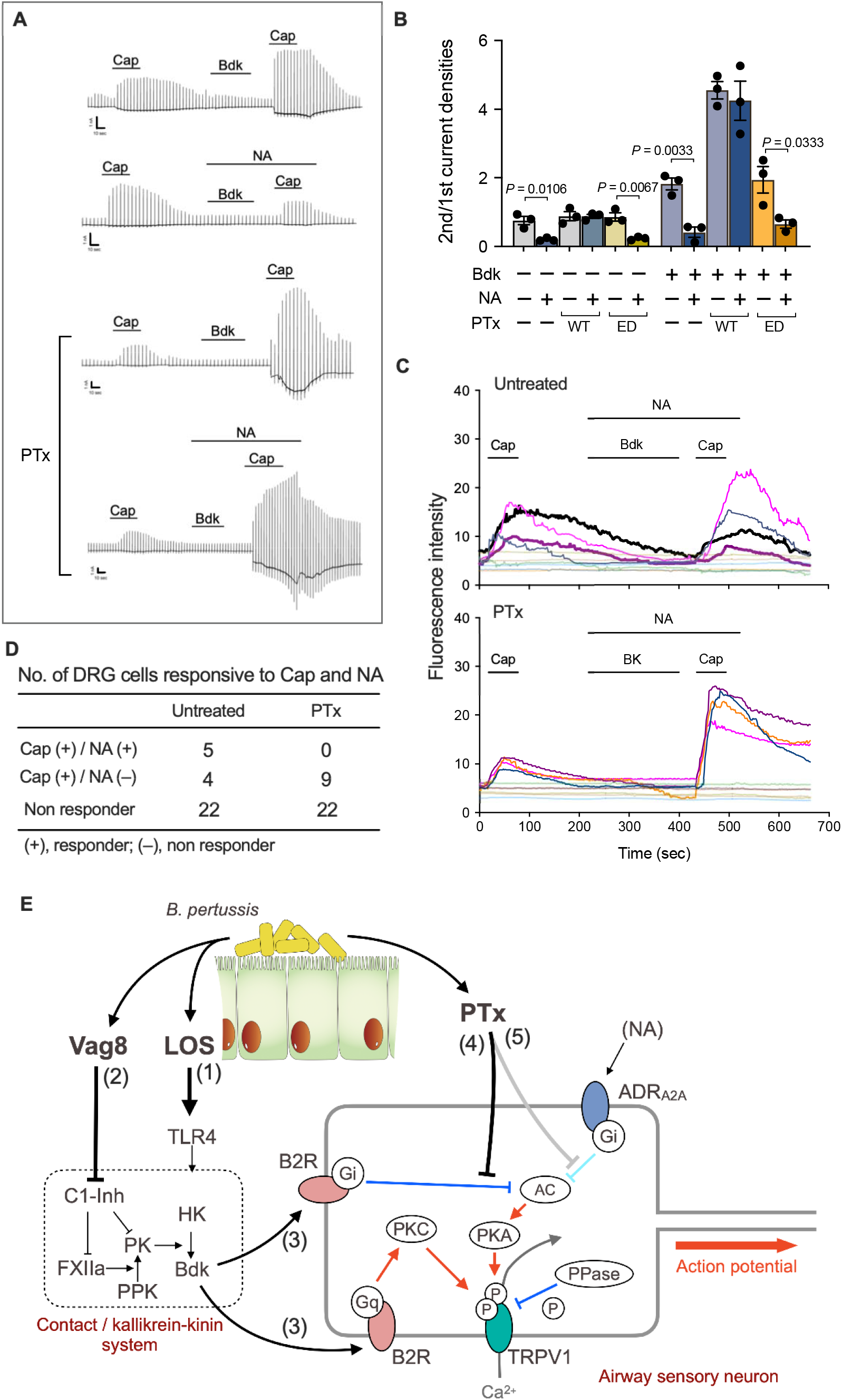
Reversal of the inhibitory effect of NA on Cap-evoked and Bdk-sensitized TRPV1 by PTx. (**A** and **B**) Current responses of HEK293T cells expressing B2R, TRPV1, and ADRA2A to repetitive applications of Cap (10 nM) and intervening applications of Bdk (100 nM) in the presence or absence of NA (100 nM). The cells were treated with 10 ng/ml PTx or PTx_ED_ or untreated prior to the whole-cell patch-clamp recording. Representative traces of Cap-evoked currents (A) and the ratio of the second peak to the initial peak of current densities induced by Cap (B) are shown. Scales at the left bottom of each trace indicate 1 nA and 10 sec on the ordinate and abscissa, respectively. Horizontal bars below the reagent names indicate the incubation period of each reagent (A). Plotted data in (B) represent means ± SEM (n = 3). (**C**) Intracellular calcium levels of DRG cells changed in response to repetitive applications of Cap (1 μM, 1 min) and an intervening application of Bdk (100 nM, 3 min) in the presence of NA (100 nM, 5 min). Temporal changes in the calcium levels are shown. Each line represents a single cell isolated from DRG, and the results from ten independent cells are depicted in each panel. The cells that responded to Cap are represented in highlight colors. The cells that responded to both Cap and NA are represented by thick lines with highlighted colors. The highlighted thin lines indicate cells responsive to Cap but irresponsive to NA. (**D**) The number of DRG cells that responded to Cap and NA. Thirty-one DRG cells were respectively examined in PTx-untreated and PTx-treated test groups. Note that no cells responded to NA in the PTx treatment group, whereas 5 out of 31 cells responded to NA in the untreated group. (**E**) The action of LOS, Vag8, and PTx to produce coughing. See Discussion in the main text.

The sensitization of TRPV1 by Bdk is considered to be mediated by the protein kinase C (PKC)-dependent pathway, which is activated via B2R-coupled G_q/11_ (Bhave et al., 2003; Mizumura et al., 2009; Numazaki et al., 2002). In contrast, the desensitization (tachyphylaxis) of TRPV1, which partly results from calcineurin-induced dephosphorylation of the channel, is reversed through phosphorylation by the cAMP-dependent protein kinase (PKA) (Bhave et al., 2002; Mohapatra, 2003; Mohapatra and Nau, 2005) (Fig. 6E). We confirmed that the Bdk-induced sensitization of TRPV1 was inhibited by calphostin C (CalpC), a PKC inhibitor, but not by H-89, a PKA inhibitor (Fig. 5C and S5B). Exacerbation of the Bdk-induced sensitization of TRPV1 by PTx was prevented by both inhibitors (Fig. 5C and S5C). An additional inhibitory effect was observed when CalpC and H-89 were simultaneously applied (Fig. 5C). Considering that the PKC inhibitor reduced the Bdk-induced sensitization of TRPV1 and that PTx inactivates G_i_ GTPases, leading to PKA activation through an increase in the intracellular cAMP level (Katada, 2012), we concluded that H-89 is likely to inhibit PTx-related events. To verify that PTx exacerbated the Bdk-induced sensitization of TRPV1 through the phosphorylation of TRPV1 by PKA, we examined HEK293T cells expressing B2R and a PKA-insensitive mutant of TRPV1 whose Ser^117^ and Thr^371^, which are phosphorylated by PKA (Bhave et al., 2002; Mohapatra, 2003; Mohapatra and Nau, 2005), were substituted by Ala (Fig. 5B and S5A). In these cells, the effects of PTx were abrogated. These results indicate that PTx exacerbates the Bdk-induced sensitization of TRPV1 through an increase in intracellular cAMP levels, followed by PKA-mediated phosphorylation of TRPV1. B2R has been reported to couple with G_q/11_, G_i_, and G_s_ GTPases (Ewald et al., 1989; Fumi et al., 1991; Gutowski et al., 1991; Lamorte et al., 1993; Liebmann et al., 1996; Mizumura et al., 2009). Our results indicate that Bdk appears to simultaneously stimulate G_q/11_- and G_i_-dependent pathways through B2R as if it simultaneously applies the accelerator and brakes on TRPV1. We conclude that PTx reverses only the inhibitory effect of Bdk on TRPV1 by uncoupling G_i_ GTPases from B2R, thereby exacerbating the Bdk-induced sensitization mediated by G_q/11_ GTPases (Fig. 6E). The involvement of G_s_ GTPases, which stimulate adenylate cyclase and subsequently PKA, in these events is probably negligible, since H-89 did not influence the Bdk-induced sensitization in the absence of PTx.

### Another possible role of PTx in the antinociceptive system

Coughs, as well as pain sensation, are initiated by nociceptive stimuli that act on nociceptors including Aδ and C fibers of the sensory nerves, where TRP channels are located. In pain sensation, nociceptive signals are modulated by antinociceptive systems, including the descending antinociceptive system conducted by serotonergic and noradrenergic neurons (Pertovaara, 2013; Tao et al., 2019). In addition, because of the similarities between the neural processing systems for pain- and cough-related inputs, several researchers have recently provided evidence for the existence of possible neural pathways that modulate the processing of the cough reflex, as well as pain sensation (Cinelli et al., 2013; Kubin et al., 2006; McGovern et al., 2017; Mutolo et al., 2014). The modulating systems for respiratory reflexes are suggested to involve the α2-adrenergic receptor, which is coupled with Gi GTPases, the target molecule for PTx (Cinelli et al., 2013; Kubin et al., 2006). If PTx inhibits such modulating systems to suppress the cough reflex, coughing will be further exacerbated. We examined this possibility using a recently reported experimental system in which noradrenaline (NA), a ligand for the α_2_ adrenergic receptor, reduced the TRPV1 activity of DRG neurons isolated from rats (Chakraborty et al., 2017; Matsushita et al., 2018) (Fig. 6A, 6B, and S6). Consistent with previous studies (Chakraborty et al., 2017; Matsushita et al., 2018), in HEK293T cells expressing TRPV1, B2R and α_2A_ adrenergic receptor (ADR_A2A_), NA inhibited the Cap-stimulated TRPV1 activity; this inhibitory effect was cancelled by pretreatment of the cells with PTx, but not with PTx_ED_ (Fig. S6). In addition, NA also inhibited the Bdk-induced sensitization of TRPV1 action and that this inhibitory effect of NA was reversed by pretreatment with PTx (Fig. 6A and 6B). Consistent results were obtained using the calcium imaging technique with DRG neurons (Fig. 6C and 6D).

## Discussion

Here, we present a series of evidence showing that LOS, Vag8, and PTx of *B. pertussis* cooperatively function to produce coughing in mice (Fig. 6E). (1) LOS stimulates Bdk generation by the kallikrein–kinin system through interaction with TLR4. (2) Vag8 accelerates Bdk generation by inhibiting C1-Inh, which is the major negative regulator of the contact system/kallikrein–kinin system. (3) Bdk sensitizes TRPV1 action, which is regulated by the phosphorylation states mediated by both PKC and PKA; phosphatases (PPase) such as calcineurin desensitize TRPV1 through dephosphorylation. Sensitization of TRPV1 by Bdk depends on B2R–G_q_-mediated PKC activation. Additionally, Bdk stimulates the inhibitory pathway of TRPV1 through B2R–G_i_-mediated inhibition of adenylate cyclase (AC) and subsequently PKA. (4) PTx reverses the B2R–Gi-mediated inhibition and exacerbates B2R–G_q_-mediated sensitization of TRPV1. Consequently, TRPV1 remains in the sensitized state and readily increase nervous excitation to evoke coughing. The sensitized state of TRPV1 continues until G_i_, which is ADP-ribosylated by PTx, is replaced by intact ones. (5) In addition, it is possible that PTx inhibits predicted negative regulation systems for the cough reflex that are relayed by G_i_ GTPase-coupled receptors, such as NA-stimulated α_2_ adrenergic receptors. The negative regulation systems for the cough reflex are yet to be clearly defined; however, step 5 deserves consideration because it may explain the long-lasting coughing in pertussis.

The causative agent of pertussis cough has long been subject to debate. One theory postulated an unidentified “cough toxin,” which is shared by the classical *Bordetella, B. pertussis, B. parapertussis*, and *B. bronchiseptica*. This idea originated from the fact that infections with the classical *Bordetella* commonly exhibit characteristic paroxysmal coughing in host animals, including humans. According to this idea, the “cough toxin” is not PTx, which is not produced by *B. parapertussis* or *B. bronchiseptica* (Cherry and Paddock, 2014). In contrast, another theory proposed that PTx significantly contributes to pertussis cough, which is supported by previous observations that rats and baboons that are experimentally infected with PTx-deficient *B. pertussis* do not exhibit coughing (Parton et al., 1994) and that immunization with pertussis toxoid protected animals from coughing but not from bacterial colonization (Hall et al., 1998; Warfel et al., 2013); these results are reaffirmed by the present study. Our results demonstrate that PTx is necessary but not sufficient to cause coughing. LOS (lipid A) as the ligand for TLR4 is common among gram-negative bacteria. Vag8 of *B. pertussis* is 97.3% identical to Vag8 of *B. parapertussis* and *B. bronchiseptica*, which themselves are completely identical. Therefore, lipid A and Vag8 are possibly the “cough toxins” shared by the classical *Bordetella. B. parapertussis* and *B. bronchiseptica*, but not *B. pertussis*, may produce another virulence factor that corresponds to PTx, which modulates the activity of the ion channels that evoke action potentials for the cough reflex. Identification of such a factor may provide an insight regarding the mechanism of cough production by the classical *Bordetella* other than *B. pertussis*.

Cough production was markedly reduced but not completely eliminated in *Tlr4*^-/-^, Kng1^-/-^, or *Trpv1*^-/-^ mice, indicating that the TLR4–Bdk–TRPV1 pathway is not the only pathway that evokes pertussis cough. LOS (lipid A) is essential for cough production. We demonstrated that lipid A-bound TLR4 slightly induced inflammatory responses; however, the mechanism through which it triggers Bdk generation remains unknown. *Tlr4*^-/-^ mice still exhibited slight coughing in response to LOS–PTx–Vag8, probably because LOS marginally induces Bdk or other cough mediators in a TLR4-independent manner. The former possibility is supported by previous studies demonstrating that lipopolysaccharide and lipid A induced Bdk generation in a mixture of FXII, PPK, and HK (Kalter et al., 1983; Roeise et al., 1988). The latter possibility is supported by the present study, which demonstrated that *Kng1*^-/-^ mice still exhibited slight coughing in response to inoculation with the bacteria or bacterial components. In addition to TRPV1, other ion channels, including TRPV4, TRPA1, TRPM8, and the purinergic P2X3 receptor are reportedly involved in signals that evoke the cough reflex (Bonvini and Belvisi, 2017). Additionally, arachidonic acid metabolites, such as prostaglandins and hydroxyeicosatetraenoic acids, and tachykinins including neurokinins and substance P are known to mediate the activation of TRP ion channels (Bonvini and Belvisi, 2017; Carr et al., 2003; Choi and Hwang, 2018; Geppetti et al., 2009; Grace et al., 2012; 2011; Ishiura et al., 2009; Kumar et al., 2017; Maher et al., 2012; 2011; Mizumura et al., 2009; Moriyama et al., 2005; Shin et al., 2002). In the present study, we negated the involvement of TRPV4, TRPA1, neurokinin receptors, and the prostaglandin E2 receptor EP3 through experiments using *Trpa1*^-/-^ mice and antagonists against these ion channels and receptors. However, the cough reflex observed in various diseases is evoked via a network or crosstalk of events involving mediators and ion channels that are not fully understood. Furthermore, the physiology of the cough reflex is not necessarily identical among animal species (Canning, 2008; Mackenzie et al., 2004). Thus, further work is required for understanding the complete mechanism of pertussis cough and applying our results to human cases.

## Acknowledgments

We thank K. Kamachi for the clinical strains of *B. pertussis*, K. Minamisawa for pRK2013, and A. Abe for pABB-CR2-Gm. We additionally acknowledge E. Mekada for critically reviewing the manuscript, and E. Hosoyamada, N. Sugi, and N. Ishihara for technical and secretarial assistance.

## Author contributions

Yu.H., K.S., and N.O. performed the main experiments. T.N., H.I., and J.K. analyzed mouse coughing using a plethysmograph. T.S., S.A., M.I., and M.T. were involved in the generation of genetically modified mice and in designing the mouse experiments. D.S. performed the electrophysiological study. Yu.H., M.T., and Ya.H. outlined the study and wrote the manuscript with contributions from the other authors.

