## Supplemental Methods and Figures for "Lipooligosaccharide, Vag8, and pertussis toxin of *Bordetella pertussis* cooperatively cause coughing in mice"

**Materials and Methods**

Bacterial strains and culture conditions

*B. pertussis* strains 18323 (Park et al., 2012) and Tohama (Parkhill et al., 2003) were maintained in our laboratory. *B. pertussis* clinical strains BP140, BP141, BP142, and BP144 were provided by K. Kamachi (National Institute of Infectious Diseases). *B. pertussis* was grown on Bordet-Gengou agar (Becton Dickinson) plates containing 1% HIPOLYPEPTON (Nihon Pharmaceutical), 1% glycerol, 15% defibrinated horse blood, and 10 μg/ml ceftibuten (BG plate). The bacteria recovered from colonies on BG plates were suspended in Strainer-Scholte (SS) medium (Stainer and Scholte, 1970) to make an OD_650_ value of 0.2 and incubated at 37°C for 12-14 h with shaking. The obtained bacteria were used as Bvg^+^-phase bacteria. Unless otherwise specified, *B. pertussis* in the Bvg^-^-phase were obtained by cultivation in the presence of 40 mM MgSO_4_. The number of colony forming units (CFU) was estimated from the OD_650_ values of fresh cultures according to the following equation: 1 OD_650_ = 3.3 × 10^9^ CFU/ml. *Escherichia coli* was grown with Luria-Bertani (LB) agar or broth. *E. coli* strains DH5α λ*pir* and HB101 harboring pRK2013 (Figurski and Helinski, 1979) were provided by K. Minamisawa (Tohoku University). The growth media were supplemented with antibiotics when necessary at the following concentrations: ampicillin, 50 μg/ml; gentamicin 10 μg/ml; kanamycin 50 μg/ml.

Construction of bacterial mutant strains

Mutant strains derived from *B. pertussis* strains 18323 and Tohama were constructed by double-crossover homologous recombination as described previously (Hiramatsu et al., 2020; Nishikawa et al., 2016). The primers and plasmids used in this study are listed in Tables S1 and S2. For the generation of 18323-∆*ptx,* -∆*fhaB*, -∆*vag8*, -∆*brkA*, and -∆*bspR* and Tohama-∆*ptx*, ~1 kbp DNA fragments of the up- and down-stream regions of the *ptx* operon and *fhaB*, *vag8*, and *brkA* genes were amplified by PCR using genomic DNA from *B. pertussis* 18323 as the template with primers ptx-U-S and ptx-U-AS, ptx-D-S and ptx-D-AS, fhaB-U-S and fhaB-U-AS, fhaB-D-S and fhaB-D-AS, vag8-U-S and vag8-U-AS, vag8-D-S and vag8-D-AS, brkA-U-S and brkA-U-AS, and brkA-D-S and brkA-D-AS, respectively. An ~0.7 kbp DNA fragment of the chloramphenicol resistant gene (*Cm^r^*) was also amplified by PCR using pKK232-8 (Addgene) as the template with primers CmR-S and CmR-AS. The PCR products of the up- and down-stream regions of each gene were ligated to the 5’ and 3’ends of the amplified *Cm^r^*, respectively, and the resultant fragments containing *Cm^r^* were inserted into the *Sma*I site of pABB-CRS2-Gm (Sekiya et al., 2001), which was provided by A. Abe (Kitasato University), using an In-Fusion HD Cloning Kit (TaKaRa Bio). The resultant plasmids (∆*ptx*-, ∆*fhaB*-, ∆*vag8*-, and ∆*brkA*-pABB-CRS2-Gm) and ∆*bspR*-pABB-CRS2-Gm (Nakamura et al., 2019) were introduced into *E. coli* DH5α λ*pir* and transconjugated into *B. pertussis* strains 18323 and Tohama by triparental conjugation with a helper strain *E. coli* HB101 harboring pRK2013. The *B. pertussis* mutant strains were isolated after confirming the replacements of the genes with *Cm^r^* by appropriate PCRs followed by agarose electrophoresis.

*B. pertussis* 18323 producing an enzymatically-inactive PTx (PTx_ED_) was constructed by site-directed mutagenesis on the gene for PTx S1 subunit (*ptxA*) to replace Arg^9^ and Glu^129^ with Lys^9^ and Gly^129^ (Buasri et al., 2012). Three distinct DNA fragments of 1.1, 0.3, and 0.9 kbp composing *ptxA* with the mutations were amplified by PCR using the genomic DNA of *B. pertussis* 18323 as the template with the combinations of primers, ptxA-S and ptxA-R9K-AS, ptxA-R9K-E129G-S and ptxA-E129G-AS, and ptxA-E129G-S and ptxA-AS, respectively. The PCR products were ligated to each other and inserted into the *Sma*I site of pABB-CR2-Gm using the In-Fusion HD Cloning Kit. The resultant plasmid, *ptxA* (R9K/E129G)-pABB-CRS2-Gm, was introduced into *E. coli* DH5α λ*pir* and transconjugated into *B. pertussis* 18323 by triparental conjugation. The resultant *B. pertussis* mutant strain was designated 18323-*ptxA* (R9K/E129G).

Mice

Three- or 6- to 10-week-old male C57BL/6J, C57BL/6N, or BALB/c mice were used (CLEA Japan and Japan SLC). *Tlr4*^-/-^ mice (Hoshino et al., 1999) were purchased from Oriental Bio Service and wild type C57BL/6J mice were used as the control. *Trpv1*^-/-^, *Trpa1*^-/-^, and *Trpv1*^-/-^/*Trpa1*^-/-^ mice were generated as previously reported (Bautista et al., 2006; Caterina et al., 2000; Kittaka et al., 2017) and the original strains were backcrossed with C57BL/6N mice (Maruyama et al., 2017). Wild-type C57BL/6N mice were used as the control. *Kng1*^-/-^ mice were generated by deleting the full length of *Kng1* gene as described previously (Abbasi et al., 2018). In brief, two-pronuclear C57BL/6J eggs were electroporated with ordered CRISPR RNAs (crRNAs; Sigma-Aldrich), trans-activating CRISPR RNA (tracrRNA; Sigma-Aldrich), and Cas9 nucleoprotein (Thermo Fisher Scientific) complexes using a NEPA21 super electroporator (NEPA GENE). The guide RNA target sequences for the 5’- and 3’-regions of *Kng1* gene were 5’-GACCTCAGGAATCTAAATAG-3’ and 5’-ACAGAGGCACGGGTGCCACA-3’, respectively. The treated zygotes were then cultured to the two-cell stage and transplanted into the oviducts of 0.5-day pseudopregnant ICR females. The founder generation was obtained by natural delivery or caesarean section. The pups obtained were genotyped by PCR using the primers Kng1-check-S1, Kng1-check-S2, and Kng1-check-AS (Table S1), and then subsequently confirmed by Sanger sequencing. The lack of Kng1 protein including HK in murine plasma was confirmed by immunoblotting with rabbit anti-Kng1 (Sigma-Aldrich) and goat anti-rabbit IgG-horseradish peroxidase (HRP) (Jackson ImmunoResearch). Wild-type littermates (C57BL/6J) were used as the control of *Kng1*^-/-^ mice. The *Kng1*^-/-^ mouse line was deposited to the RIKEN BioResource Research Center and Center for Animal Resources and Development, Kumamoto University. All animal experiments were approved by the Animal Care and Use Committee of the Research Institute for Microbial Disease, Osaka University and carried out according to the Regulations on Animal Experiments at Osaka University.

Cough analysis

Six- to 10-week-old mice were anesthetized with a mixture of medetomidine (Kyoritsu Seiyaku), midazolam (Teva Takeda Pharma), and butorphanol (Meiji Seika Pharma) at final doses of 0.3, 2, and 5 mg/kg body weight, respectively, and intranasally inoculated with *B. pertussis* or its components in 50 μl of SS medium using a micropipette with a needle-like tip. The amount of bacteria were confirmed by counting colonies after cultivation of the inoculums on BG plates. The numbers of murine coughs from day 4 to 14 post-inoculation were enumerated as previously described with slight modifications (Nakamura et al., 2019). The mice were isolated individually in disposable clear plastic cages, which were laid on a sound-proof sheet and covered with plastic cardboard. Coughing was recorded by a video camera (Canon HD ivisHF21, Canon) equipped with two monaural microphones (AT9903, Audio-Technica Co.) every day for 5 min a day. The recorded data were processed with Adobe Premier Pro CS5.5 (Adobe Systems Incorporated) on a computer and displayed as movies along with sound waveforms. Coughs were checked by characteristic waveforms and the coughing postures of mice, and enumerated by an observer who did not know detailed information about the experiments. After 14 days of post-inoculation, the mice were euthanized with pentobarbital and their tracheas and lungs were aseptically excised, minced and homogenized in Dulbecco’s phosphate buffered saline (PBS) with a BioMasher (Nippi) and a Polytron PT1200E (Kinematica), respectively. The resultant tissue extracts were serially diluted with PBS and spread on BG plates. The bacteria on the plates were cultivated at 37°C for 3-4 days, and the numbers of CFU were enumerated.

For the preparation of bacterial lysates, *B. pertussis* cultivated in SS medium were collected by centrifugation at 8,000 × *g* for 10 min. The bacteria resuspended in PBS were disrupted by 5 rounds of 2-min sonication with Bioruptor (Cosmo Bio). The sonicated suspensions were centrifuged at 12,000 × *g* for 5 min, and the cell lysates were obtained after filtration of the supernatants through a 0.22-μm filter (Millex GV, Merck Millipore). The obtained cell lysates were intranasally inoculated daily for 5 days at 50 μg/50 μl into mice that were anesthetized with isoflurane using an anesthetizer (MK-A110; Muromachi Kikai). LOS, Vag8, and PTx were inoculated in a similar way to the bacterial cell lysate at 4 × 10^4^ EU, 500 ng, and 200 ng in 50 µl, respectively. Antagonists of B1R (Des-Arg^9^-[Leu^8^]-Bradykinin; PEPTIDE Institute), B2R (Icatibant; PEPTIDE Institute), TRPV1 (Capsazepine; Wako Pure Chemical Industries), TRPA1 (HC-030031; Wako Pure Chemical Industries), TRPV4 (HC-067047; Sigma-Aldrich), NK1 (Spantide; PEPTIDE Institute), NK2 (GR159897; R&D Systems), NK3 (SB222200; Sigma-Aldrich), and EP3 (L-798106; Sigma-Aldrich) were intraperitoneally inoculated at 20 or 60 nmol/300 μl daily for 5 days into mice immediately before intranasal inoculation of the *B. pertussis* cell lysates or PTx, LOS, and Vag8. The numbers of murine coughs from day 4 to 14 after the initial inoculation were enumerated as described above.

Vaccination

Three-week-old male C57BL/6J mice were intraperitoneally injected with 1/4, 1/10 and 1/25 the human dose of an adsorbed diphtheria-purified pertussis-tetanus inactivated polio combined vaccine (DPT-IPV) (TETRABIK, BIKEN, Osaka, Japan) or 4 mg/100 µl of aluminum hydroxide (Imject Alum; Thermo Fisher Scientific) as the negative control. Booster injections were performed three weeks after the primary injection. The mice were bled just before the primary injection and seven days after the booster injection, and anti-PTx antibody titers in the sera were estimated by ELISA as follows. Ninety-six-well polystyrene plates (ELISA Plate H; Sumitomo Bakelite) were coated with 0.1 μg/well of PTx at 4°C overnight and then blocked with 300 μl/well of PBS containing 0.5% BSA at 37°C for 1 h. Serum samples were ten-fold serially diluted in PBS, added to the wells of the plates, and incubated at 37°C for 1 h. The anti-PTx antibodies in the serum were sequentially probed with goat anti-mouse IgG-HRP (Jackson ImmuneResearch) at 37°C for 1 h. After each step, the wells were washed five times with PBS containing 0.05% Tween 20. One hundred microliters of 0.1% TMBZ (3, 3’, 5, 5’-tetramethylbenzidine; Dojindo Laboratories) in 0.1 M citrate-acetate buffer, pH 6.0, containing 0.01% H_2_O_2_ was added to each well and allowed to react at room temperature for 30 min. The reactions were stopped by 100 μl/well of 1 M H_2_SO_4_. The optical density of each well was read on a Multi-Detection Microplate Reader at 450 nm. After seven days of the booster immunization, the mice were intranasally inoculated with *B. pertussis* (5 × 10^6^ CFU), and then the numbers of murine coughs were enumerated as described above. The colonization levels in murine tracheas and lungs were assessed after 14 days of the bacterial inoculation as described above.

Recombinant Vag8s

The expression vectors for HAT-tagged recombinant Vag8 proteins, pCold II-HAT-*vag8*_Tohama_ and pCold II-HAT-*vag8*_18323_, pCold II-HAT-*vag8*_102-596_, _102-548, and_ _102-479_, which were constructed as previously described (Onoda et al., 2020), were introduced into *E. coli* BL21 (DE3). The Vag8 proteins were expressed by incubation at 15°C for 24 h in the presence of 1 mM isopropyl-β-D-thiogalactopyranoside (IPTG). The bacteria were collected by centrifugation and disrupted by sonication in 50 mM sodium phosphate buffer, 300 mM NaCl, pH 8.0 (Buffer A), containing 10 mM imidazole. After centrifugation, the supernatants were independently applied to a column of HIS-Select Nickel Affinity Gel (Sigma-Aldrich) equilibrated with Buffer A containing 10 mM imidazole. After nonabsorbed substances had been washed out of the column with Buffer A containing 10 mM imidazole, the recombinant Vag8 proteins were eluted with Buffer A containing 300 mM imidazole. Imidazole in the Vag8 fraction was removed by dialysis against PBS.

LOS preparation

LOS was purified from *B. pertussis* strains 18323 and Tohama by the hot-phenol method (Minnick, 1994). The purity of the LOS preparation was assessed by sodium dodecyl sulfate-polyacrylamide gel electrophoresis (SDS-PAGE; 20% gel), followed by silver staining using Sil-Best Stain One (Nacalai tesque). The amount of endotoxin was measured by the Limulus Color KY Test (Wako Pure Chemical Industries) and expressed as endotoxin units (EU) according to the manufacturer’s instruction. In an independent experiment, endotoxin in *B. pertussis* cell lysates was removed with Detoxi-Gel Endotoxin Removing Columns (Thermo Fisher Scientific).

Measurement of bradykinin and cytokine levels in BALF

Six- to 8-week-old wild type or *Tlr4*^-/-^ mice were intranasally inoculated daily for 5 days with PTx, LOS, and/or Vag8. On the indicated days after the first inoculation, the mice were euthanized with pentobarbital, and their tracheas were exposed by incising the necks in the median line. A catheter (Surflo Flash 20G; Terumo) was inserted into the exposed tracheas, 0.5 ml of PBS was slowly injected, and bronchoalveolar lavage fluid (BALF) was thoroughly aspirated. The BALF was centrifuged at 12,000 × *g* for 5 min, and the supernatant was collected. The concentrations of bradykinin and cytokines in the supernatant were measured by Mouse Bradykinin ELISA Kit (MyBioSource), Mouse IL-1β/IL-1F2 DuoSet ELISA (R&D Systems, Code: DY401-05), Mouse IL-6 DuoSet ELISA (R&D Systems, Code: DY406-05), and Mouse TNF-α DuoSet ELISA (R&D Systems, Code: DY410-05).

Electrophysiology

HEK293T cells were maintained in Dlbecco’s modified Eagle medium (DMEM; Sigma-Aldrich or Wako) supplemented with 10% fetal bovine serum (FBS; Biowest) at 37°C under 5% CO_2_ in air. The cells that were seeded on a 35-mm dish (IWAKI) at 5 × 10^5^ cells/well and grown overnight were transfected with pRc/CMV-*mB2R* (Wang et al., 2008), pcDNA-*hTRPV1* (Wang et al., 2008), pcDNA-*hTRPV1_S117A/T371A_*, pCMV-SPORT6-*hADRA2A* (DNAFORM, ID: 6198830), and/or pGreen-Lantern 1 (Li et al., 2019), which is a humanized GFP expression vector, using Lipofectamine reagent (Thermo Fisher Scientific) according to the manufacturer’s instructions. pcDNA-*hTRPV1_S117A/T371A_* was generated from pcDNA-*hTRPV1* by site-directed mutagenesis to replace Ser^117^ and Thr^371^ with Ala using a PrimeSTAR Mutagenesis Basal Kit (TaKaRa Bio) with the primer sets S117A-S and S117A-AS, and T371A-S and T371-AS (Table S1). After incubation for 3-4 h, the cells were reseeded on 12-mm coverslips (Matsunami Glass) and further incubated for 2 days. Whole-cell patch-clamp recordings were performed in a GFP-positive single cell which had been treated with PTx or PTx_ED_ at 10 ng/ml for 24-30 h prior to the recordings, as previously described with slight modifications (Sugiura et al., 2002). The standard bath solution (140 mM NaCl, 5 mM KCl, 2 mM MgCl_2_, 2 mM CaCl_2_, 10 mM Hepes, and 10 mM glucose, pH7.4) was replaced with Ca^2+^-free bath solution, in which CaCl_2_ of the standard bath solution was replaced with 5 mM EGTA, immediately prior to the recording. Electrodes were filled with the pipette solution (140 mM KCl, 5 mM EGTA, 10 mM Hepes, pH 7.4). Currents generated across the cells, which were treated with 10 nM capsaicin (Nacalai tesque), 100 nM bradykinin (PEPTIDE Institute), 100 nM noradrenaline (Sigma-Aldrich), 1 µM calphostin C (Merck Millipore), and/or 1 µM H-89 (Merck Millipore) for arbitrary periods of time, were recorded with an Axopatch 200B amplifier (Molecular Devices), filtered at 5 kHz with a low-pass filter, and digitized with Digidata 1440A (Axon Instruments). The membrane potential was clamped at -60 mV and voltage ramp-pulses from -100 to + 100 mV (0.5 sec) were applied every 5 sec. Data were acquired with pCLAMP 10 (Axon Instruments), and the plots were generated with OriginPro 2016 (OriginLab). Current densities (pA/pF) of the peak currents induced by the first and second capsaicin treatments were calculated as the quotient of the current amplitude (pA) divided by whole cell capacitance (pF).

Calcium imaging

DRG cells were isolated from 7-week-old male C57BL/6J mice that were euthanized with pentobarbital as described previously (Kittaka et al., 2017). The DRG cells were suspended in 0.5 ml/mouse of Eargle’s balanced salts solution containing 10% FBS, 50 units/ml penicillin, 50 µg/ml streptomycin, 1% GlutaMAX Supplement (Thermo Fisher Scientific), and MEM vitamin solution (Sigma Aldrich), seeded on a 35-mm glass base dish, and incubated at 37°C under 5% CO_2_ in air for 24-30 h with or without PTx or PTx_ED_. For the calcium imaging, DRG cells were incubated with Hank’s Balanced Salt solution (Sigma-Aldrich), 20 mM Hepes, pH7.4 (HBSS-Hepes) containing 4 µM Fluo-4 AM (Dojindo), 1.25 mM probenecid, and 0.04% cremophor EL at room temperature for 30 min, and then washed three times with HBSS-Hepes to remove extracellular Fluo-4 AM. Fluorescence images of the Fluo-4-loaded cells, which were treated with 1 µM capsaicin, 100 nM bradykinin, 100 nM noradrenaline, 1 µM calphostin C, and/or 1 µM H-89 at the appropriate periods, were captured at 2-sec intervals using a fluorescence microscope (Olympus BX51). The relative fluorescence intensities of independent cells in each fluorescence image were quantified with Metamorph 7.6 (Molecular Devices). The increased transient peak values of the fluorescence intensity induced by the first and second capsaicin treatments were calculated by subtracting the fluorescence value obtained immediately before the initial capsaicin treatment from that of the maximum fluorescence peak.

Statistical analysis

Unless otherwise specified, one-way analysis of variance (ANOVA) with Dunnett’s or Tukey’s multiple-comparison test was performed to evaluate differences between test groups by using Prizm 8 (GraphPad Software).

Others

PTx was purified from culture supernatants of *B. pertussis* strains 18323 (wild-type and *ptx*_R9K/E129G_ mutant) and Tohama as previously reported (Skelton and Wong, 1990). PTx was intranasally inoculated daily for 5 days at 0.2 μg/50 μl into mice. The ADP-ribosylation of G_i_ proteins by PTx in HEK293T cells and DRG cells were confirmed by immunoblotting of the cell lysates with rabbit anti-poly/mono ADP-ribose antibody (Cell Signaling Technology), mouse anti-G_αi-1_ (Santa Cruz Biotechnology), or mouse anti-GAPDH (Wako Pure Chemical Industries) antibodies followed by goat anti-rabbit IgG-HRP or goat anti-mouse IgG-HRP (Jackson ImmunoResearch).The protein concentration of the test materials used in this study were measured using Micro BCA Protein Assay Kit (Thermo Fisher Scientific). Synthesized lipid A of *E. coli* (code no. 24005-s) was purchased from PEPTIDE Institute Inc. In immunoblotting, the target proteins were visualized by enhanced chemiluminescence using an Immobilon Western (Merck Millipore) and LAS-4000mini Luminescent Image Analyzer (GE Healthcare) or Amersham Imager 600 UV (GE Healthcare).

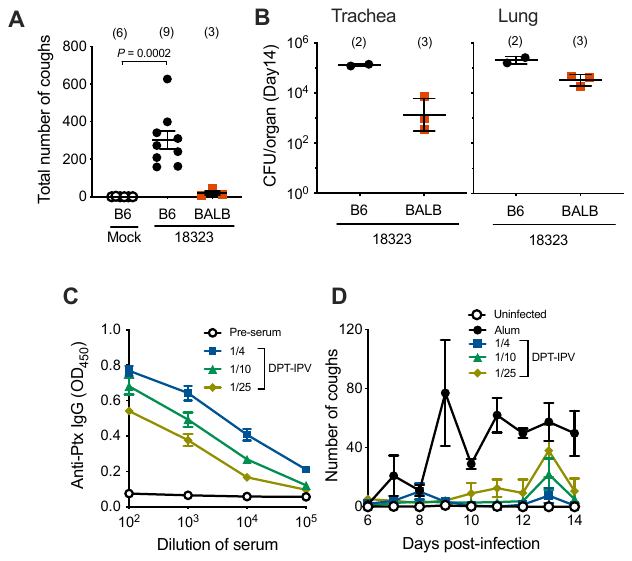

**Fig. S1. Differences in the cough response between C57BL/6J and BALB/c mice and effects of immunization with DPT-IPV**

(**A** and **B**) Comparison of C57BL/6J (B6) and BALB/c (BALB) mice in cough production after inoculation with *B. pertussis* 18323. Mice were intranasally inoculated with *B. pertussis* 18323 as described in Materials and Methods, and the numbers of coughs were counted for 9 days from days 6 to 14 post-inoculation of the bacteria (A). The number of bacteria recovered from the tracheas and lungs was counted on day 14 (B). Each horizontal bar represents the mean ± SEM (A) or geometric mean ± SD (B). (**C** and **D**) Cough production in DPT-IPV-immunized mice. Mice that were immunized with 1/4, 1/10, or 1/25 the human dose of the DPT-IPV (n = 5 for each test group) were intranasally inoculated with 5 × 10^6^ CFU of *B. pertussis* 18323 or SS medium (Uninfected, n = 3) as described in Materials and Methods. The sera of mice were obtained just before (Pre-serum, n = 15) and 28 days (n = 5) after the primary immunization. The levels of anti-PTx antibody in sera of immunized mice were determined using ELISA (C). The numbers of coughs were counted for 9 days from days 6 to 14 post-inoculation of the bacteria (D). For control, aluminum hydroxide adjuvant without the immunogen was injected (Alum, n = 4). Each plot represents the mean ± SEM.

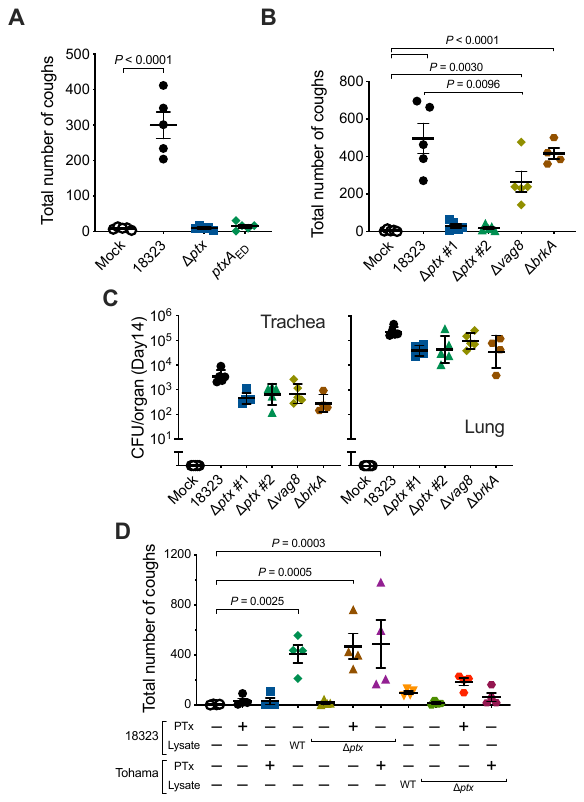

**Fig. S2. Involvement of PTx and Vag8 in *B. pertussis*-induced coughing.**

(**A**-**C**) Cough production in mice inoculated with bacterial cell lysates or living bacteria. Mice were intranasally inoculated with 50 µg of cell lysates of *B. pertussis* 18323 wild-type, Δ*ptx*, *ptxA*_ED_ or PBS (Mock, 50 µl) (A) or with *B. pertussis* 18323 wild-type, two distinct clones of Δ*ptx*, Δ*vag8*, or Δ*brkA* or SS medium (Mock, 50 µl) (B). The numbers of coughs were counted for 5 min/mouse/day from day 6 to 14 post-inoculation (A and B). The number of bacteria recovered from the tracheas and lungs on day 14 post-inoculation was enumerated (C). (**D**) Cough-inducing activities of the cell lysates and PTx from the *B. pertussis* strains 18323 and Tohama. Mice were intranasally inoculated with various combinations of 50 µg of cell lysates and/or 200 ng of PTx of *B. pertussis* strains 18323 or Tohama (wild-type or Δ*ptx*) in 50 µl of PBS. The number of coughs in the mice was counted as described above, and the total number of coughs per mouse from days 6 to 14 is presented. Each horizonal bar represents the mean ± SEM (A and B, n = 5; D, n=4) or geometric mean ± SD (C, n = 5).

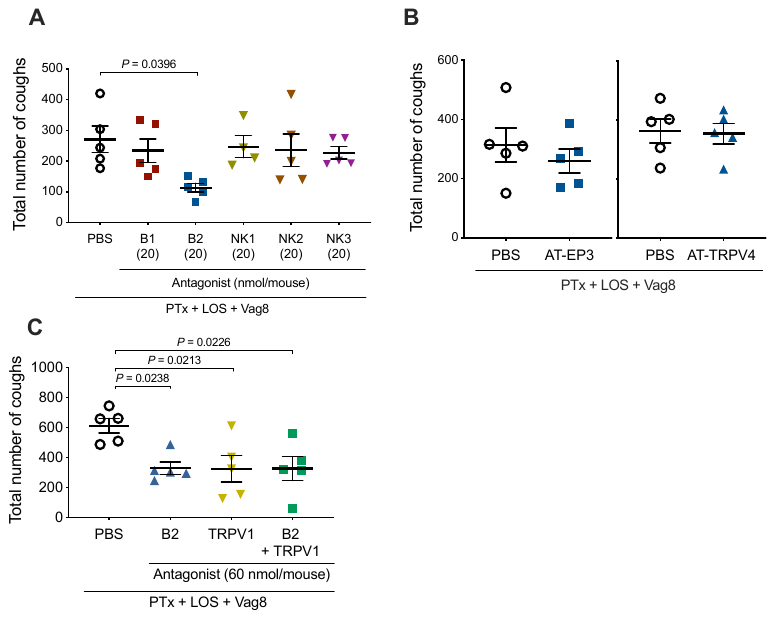

**Fig. S3. Effects of antagonists against cough reflex-related signal pathways on *B. pertussis*-induced coughing.**

Mice were intraperitoneally inoculated with 20 (A) or 60 (B and C) nmol of antagonists against Bdk receptors (B1 and B2), neurokinin receptors (NK1-3), prostaglandin E2 receptor (AT-EP3), TRPV4 (AT-TRPV4), TRPV1 or PBS (300 µl) and subsequently, intranasally inoculated with PTx (200 ng), LOS (4 × 10^4^ EU), and Vag8 (500 ng) every day from days 0 to 4. The number of coughs was counted for 5 min/mouse/day. The total number of coughs per mouse from days 6 to 14 is presented. Each horizontal bar represents the mean ± SEM (n = 5).

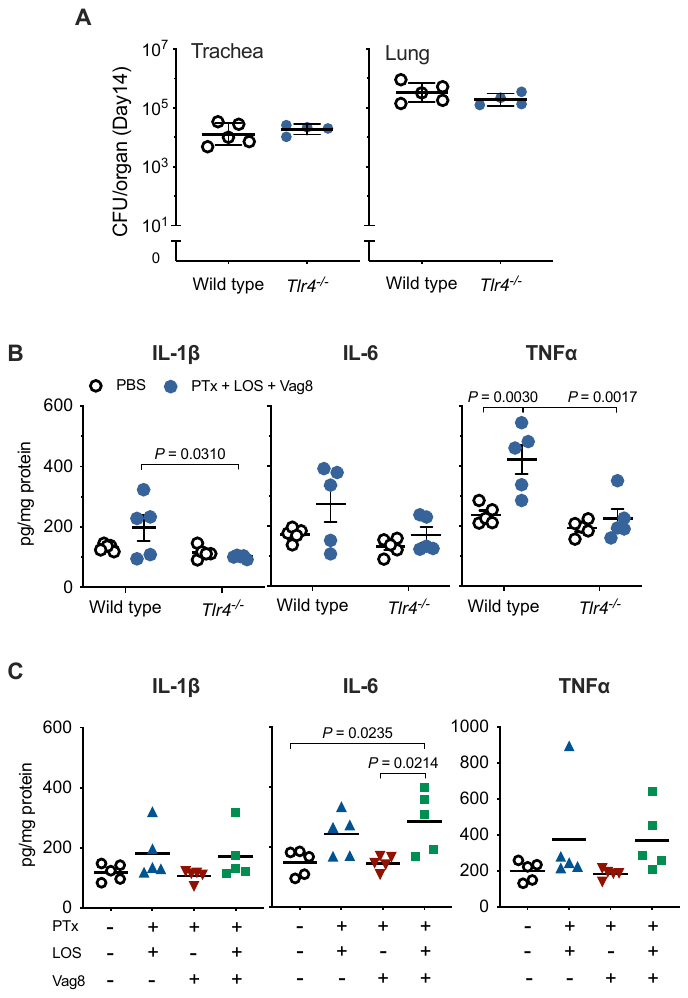

**Fig. S4. Role of LOS in *B. pertussis*-induced coughing.**

**(A)** Colonization by *B. pertussis* of *Tlr4*^-/-^ mice. Wild-type or *Tlr4*^-/-^ mice were intranasally inoculated with 5 × 10^6^ CFU of *B. pertussis* 18323, and the number of bacteria recovered from murine tracheas and lungs was counted on day14 post-inoculation. Each horizonal bar represents the geometric mean ± SD. (**B** and **C**) Concentrations of IL-1β, IL-6, and TNFα in the BALF of mice inoculated with PTx, LOS, and/or Vag8. Wild type (B and C) and *Tlr4*^−/−^ (B) were inoculated with the samples as described in Materials and Methods, and the concentrations of the cytokines in BALF on day 4 were determined by ELISA.

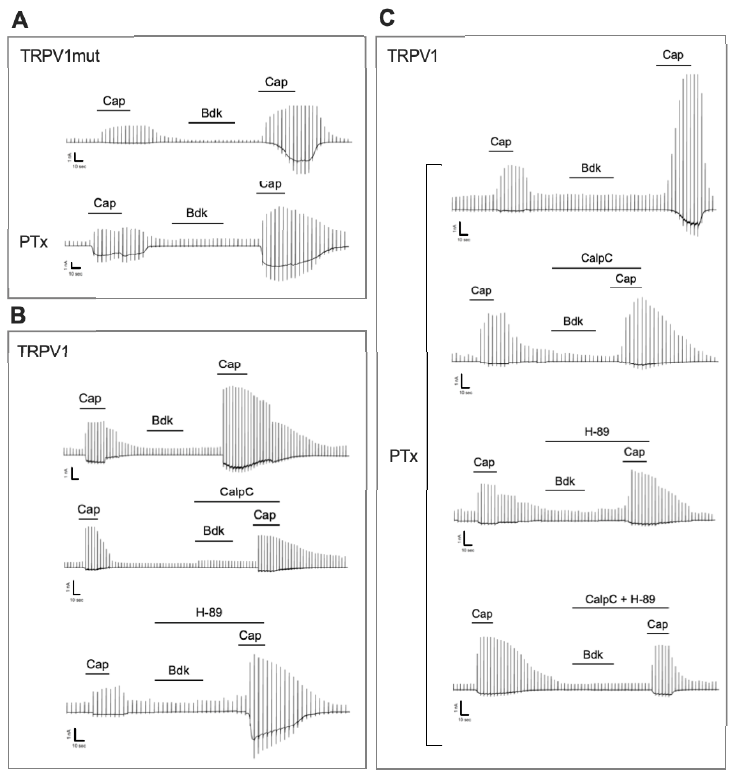

**Fig. S5. Representative traces of capsaicin-evoked currents of HEK293T cells expressing B2R and TRPV1 or TRPV1_mut_.**

(**A**) Responses of cells expressing TRPV1_mut_ to PTx treatment. (**B** and **C**) Effects of CalpC, H-89, and/or PTx on the Cap-evoked and Bdk-sensitized responses of the cells. Cells expressing mB2R and hTRPV1 (B and C) or hTRPV1_mut_ (A) were transiently treated with 10 nM capsaicin (Cap) and 100 nM Bdk in the presence or absence of CalpC (1 µM) and/or H-89 (1 µM). The cells were treated with 10 ng/ml PTx for 24–30 h prior to the whole-cell patch-clamp recordings. Horizontal bars indicate the exposure time of each reagent. Scales at the left bottom of each trace indicate 1 nA and 10 sec on the ordinate and abscissa, respectively.

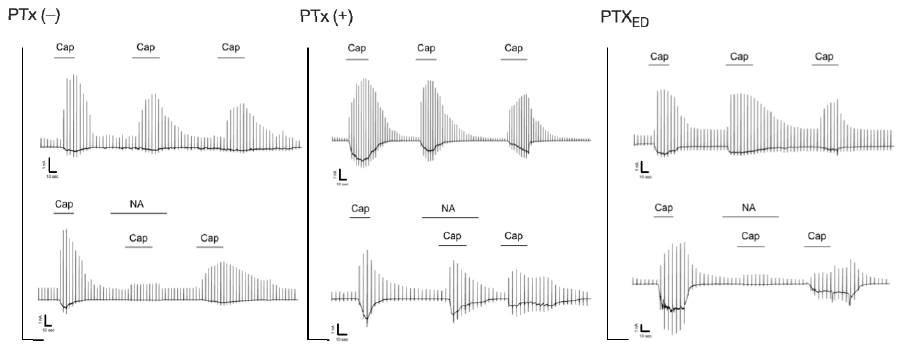

**Fig. S6. Reversal of the inhibitory effect of NA on Cap-evoked TRPV1 activity by PTx.**

Representative traces of capsaicin-induced currents recorded by the whole-cell patch-clamp technique. HEK293T cells expressing TRPV1 and ADR_A2A_ were previously treated or untreated with PTx or PTx_ED_ and stimulated by repetitive applications of capsaicin (Cap, 10 nM) in the presence or absence of NA (100 nM).

**Table. S1 Primers used in this study**

| Name | Sequence (5’ to 3’) | Template | Application |
| --- | --- | --- | --- |
| ptx-U-S | GATCCGAGCTCTCCCAACCCGTCGTGGTGCAGGA | 18323 genomic DNA | Δ*ptx-*pABB-CRS2-Gm |
| ptx-U-AS | TTAGCTTCCTTAGCTAGTGCAACGCATCCCGTC |  |  |
| ptx-D-S | CGCGCCATTTAAATGGCGTCGATATGCTGAGCC |  |  |
| ptx-D-AS | ATTTGTGGAATTCCCGTTCGTCATTGCCTGCC |  |  |
| fhaB-U-S | GATCCGAGCTCTCCCAAGAACGGCACCTGGAC | 18323 genomic DNA | Δ*fhaB*-pABB-CRS2-Gm |
| fhaB-U-AS | TTAGCTTCCTTAGCTAAATAGGTAGTCGCGGCC |  |  |
| fhaB-D-S | CGCGCCATTTAAATGATTCCGACCAGCGAAGTG |  |  |
| fhaB-D-AS | ATTTGTGGAATTCCCTCAGCTTCTGCAGCAGG |  |  |
| vag8-U-S | GATCCGAGCTCTCCCGCGCTTGGGGATTTTTCAGGTTT | 18323 genomic DNA | Δ*vag8-*pABB-CRS2-Gm |
| vag8-U-AS | TTTAGCTTCCTTAGCTGACCTGAACCACCAGCCCCTGT |  |  |
| vag8-D-S | CGCGCCATTTAAATGTATCGCTACAGCTGGTGACCGCG |  |  |
| vag8-D-AS | TTTGTGGAATTCCCGTGCTGCTGGCCGAACAGCTCTAC |  |  |
| brkA-U-S | GATCCGAGCTCTCCCCTGCGCTGAATAATAGATCCACA | 18323 genomic DNA | Δ*brkA-*pABB-CRS2-Gm |
| brkA-U-AS | TTTAGCTTCCTTAGCTGTGCCACCAAAAGAGAAGTTGA |  |  |
| brtA-D-S | CGCGCCATTTAAATGGCGAGCTCCAATGAAAAACCCCG |  |  |
| brkA-D-AS | TTTGTGGAATTCCCTACTTCATCAACAGCCCGGAAAAG |  |  |
| CmR-S | AGCTAAGGAAGCTAAAATGGAG | pKK232-8 | Cm resistant gene |
| CmR-AS | CATTTAAATGGCGCGCCTT |  |  |
| ptxA-S | GATCCGAGCTCTCCCTACACCCTGTTGCTGTCGT | 18323 genomic DNA | *ptxA* (R9K/E129G)-pABB-CRS2-Gm |
| ptxA-R9K-AS | TTTGTATACGGTGGCGGGAGGATC |  |  |
| ptxA-R9K-E129G-S | GCCACCGTATACAAATATGACTCCCGCCCG |  |  |
| ptxA-E129G-AS | GCCAGATATCCGCTCTGGTAGGTGGCCA |  |  |
| ptxA-E129G-S | CCAGAGCGGATATCTGGCACACCGG |  |  |
| ptxA-AS | ATTTGTGGAATTCCCCGCGTACGGAGATGCCGG |  |  |
| Kng1-check-S1 | CAACTCAGGCCGTGTGTTTG | *Kng1* gene | check for *Kng1* gene deletion |
| Kng1-check-S2 | GCAGAGCCATCCTGAGGAAA |  |  |
| Kng1-check-AS | ACTCCCCAGCTGCTATCAGA |  |  |
| S117A-S | CGCAGGGCTATCTTTGAAGCCGTTGCTCAGAAT | pcDNA-*hTRPV1* | pcDNA-*hTRPV1*_S117A/T371A_ |
| S117A-AS | AAAGATAGCCCTGCGATCATAGAGCCTGAGGGT |  |  |
| T371-S | GAAGTTCGCCGAGTGGGCCTACGGGCCCGTGCA |  |  |
| T371-AS | CACTCGGCGAACTTCCTGGACAGGTGCCTGCAC |  |  |

**Table. S2 Plasmids and bacterial strains used in this study**

| Plasmids and strains | | Description | Source or reference |
| --- | --- | --- | --- |
| Plasmids | |  |  |
|  | pRK2013 | Km^r^, RK-2 derivative with ColE1 replicon containing *tra*,  helper plasmid for conjugative transfer | (Figurski and Helinski, 1979) |
|  | pKK232-8 | Cm^r^, cloning vector | Addgene |
|  | pABB-CRS2-Gm | Gm^r^, R6K-derived suicide vector | (Sekiya, et.al., 2001) |
|  | Δ*ptx*-pABB-CRS2-Gm | *ptx*-operon deletion cloned into pABB-CRS2-Gm | This study |
|  | Δ*fhaB-*pABB-CRS2-Gm | *fhaB* deletion cloned into pABB-CRS2-Gm | This study |
|  | Δ*brkA-*pABB-CRS2-Gm | *brkA* deletion cloned into pABB-CRS2-Gm | This study |
|  | Δ*vag8-*pABB-CRS2-Gm | *vag8* deletion cloned into pABB-CRS2-Gm | This study |
|  | *ptxA* (R9K/E129G)-pABB-CRS2-Gm | *ptxA* (R9K/E129G) mutant cloned into pABB-CRS2-Gm | This study |
|  | pCold II-HAT-*vag8*_18323_ | pCold II-HAT carrying *vag8* derived from *B. pertussis* 18323 | (Onoda, et.al., 2020) |
|  | pCold II-HAT-*vag8*_Tohama_ | pCold II-HAT carrying *vag8* derived from *B. pertussis* Tohama | (Onoda, et.al., 2020) |
|  | pCold II-HAT-*vag8*_102-596_, _102-548, and_ _102-479_ | pCold II-HAT carrying truncated forms of *vag8* | (Onoda, et.al., 2020) |
|  | pRc/CMV-*mB2R* | pRc/CMV carrying mouse *B2R* cDNA | (Wang, et.al., 2008) |
|  | pcDNA-*hTRPV1* | pcDNA3.1 carrying human *TRPV1* cDNA | (Wang, et.al., 2008) |
|  | pcDNA-*hTRPV1*_S117A/T371A_ | pcDNA3.1 carrying cDNA of human *TRPV1*, in which Ser and Thr were replaced with Ala at amino acid position 117 and 371 | This study |
|  | pCMV-SPORT6-h*ADRA2A* | pCMV-SPORT carrying human *ADRA2A* cDNA | DNAFORM |
|  | pGreen Lantern 1 | Humanized GFP expression vector | (Li, et.al., 2019) |
| *B. pertussis* | |  |  |
|  | 18323 | Type strain | (Park, et.al., 2012) |
|  | Tohama | Vaccine strain | (Parkhill, et.al., 2003) |
|  | BP140 | Clinical strain isolated from a pertussis patient | K. Kamachi |
|  | BP141 | Clinical strain isolated from a pertussis patient | K. Kamachi |
|  | BP142 | Clinical strain isolated from a pertussis patient | K. Kamachi |
|  | BP144 | Clinical strain isolated from a pertussis patient | K. Kamachi |
|  | 18323-Δ*ptx* | 18323 derivative, Δ*ptx*::*Cm^r^* | This study |
|  | Tohama-Δ*ptx* | Tohama derivative, Δ*ptx*::*Cm^r^* | This study |
|  | 18323-Δ*fhaB* | 18323 derivative, Δ*fhaB*::*Cm^r^* | This study |
|  | 18323-Δ*brkA* | 18323 derivative, Δ*brkA*::*Cm^r^* | This study |
|  | 18323-Δ*vag8* | 18323 derivative, Δ*vag8*::*Cm^r^* | This study |
|  | 18323-Bvg^+^-lock | 18323 derivative producing BvgS, in which Arg was replaced with His at amino acid position 570 (Bvg^+^ phase-locked mutant) | (Hiramatsu, et.al., 2020) |
|  | 18323-Bvg^-^-lock (18323Δ*bvgS*) | 18323 derivative, in which BvgS was deleted from 542 to 1020 amino acid positions (Bvg^-^ phase-locked mutant) | (Hiramatsu, et.al., 2020) |
|  | 18323-*ptxA* (R9K/E129G) | 18323 derivative producing PT S1 subunit, in which Arg and Glu were replaced with Lys and Gly at amino acid positions 9 and 129 | This study |

**Movie S1.** A mouse 10 days after inoculation with SS medium.

**Movie S2**. A mouse 10 days after inoculation with 5 × 10^6^ CFU of *B. pertussis*.
